# Matrix stiffness modulates infection of endothelial cells by *Listeria monocytogenes* via expression of cell surface vimentin

**DOI:** 10.1101/198739

**Authors:** Effie E. Bastounis, Yi-Ting Yeh, Julie A. Theriot

## Abstract

Extracellular matrix (ECM) stiffness is one of many mechanical forces acting on mammalian adherent cells that influence cellular function. We have addressed the open question of how ECM stiffness might alter the susceptibility of host cells to infection by bacterial pathogens. We manufactured hydrogels of varying physiologically-relevant stiffness and seeded human microvascular endothelial cells (HMEC-1) on them. We then infected HMEC-1 with the bacterial pathogen *Listeria monocytogenes* (Lm) and found that adhesion of Lm onto host cells increases monotonically with increasing matrix stiffness, an effect that requires the activity of focal adhesion kinase (FAK). We identified cell surface vimentin as a candidate surface receptor mediating stiffness-dependent adhesion of Lm to HMEC-1, and demonstrated that bacterial infection of these host cells is decreased when surface vimentin is perturbed. Our results provide the first evidence that ECM stiffness can mediate the susceptibility of host cells to bacterial infection.

## INTRODUCTION

The extracellular environment of cells provides both chemical and mechanical stimuli to influence cell behavior and function (Geiger et al., 2009, Chien et al., 2005). Extracellular matrix stiffness (ECM), one of the many mechanical forces acting on cells, is an important determinant of cellular behavior for most adherent mammalian cells (Gattazzo et al., 2014; Wells, 2008). Cells can sense the stiffness of their matrix, which can vary over many orders of magnitude, and accordingly alter their motility, adhesion, growth and differentiation (Discher et al., 2005; Birukova et al., 2013). Yet the exact pathways whereby cells sense mechanical signals and transduce them to generate biological signal cascades and specific cellular responses are not yet fully understood (Trepat et al., 2008).

In the context of host-pathogen interactions, effects of matrix stiffness variation may be most interesting for infectious agents that have the capacity to infect many different kinds of tissues in the human body. One such agent is the food-borne facultative bacterial pathogen *Listeria monocytogenes* (Lm). After initial invasion into the intestinal epithelium, Lm is able to spread through the vasculature to distant organs, and can cause serious complications such as meningitis and late-term spontaneous abortion by virtue of its unusual ability to penetrate and cross a wide variety of endothelial barriers including the blood-brain barrier and the placenta (Jackson et al., 2010; Vazquez-Boland et al., 2001). Lm has a broad range of susceptible host animals, and uses multiple pathogenic strategies to achieve infection of a wide variety of tissues within each host. It can directly adhere to and invade intestinal epithelial cells and hepatocytes using bacterial surface proteins belonging to the internalin family, such as InlA, InlB and InlF, which interact with host cell surface receptors (Mengaud et al., 1996; Shen et al., 2000; Kirchner and Higgins, 2008). Lm can also directly adhere to and invade the vascular endothelial cells (VECs) that line the inner lumen of blood vessels, using several distinct molecular mechanisms that may vary depending on the subtype of VEC being infected (Drevets et al., 1995; Greiffenberg et al., 1997; Wilson and Drevets, 1998; Greiffenberg et al., 1998; Parida et al., 1998; Rengarajan et al., 2016).

Vascular endothelial cells (VECs) are known to be highly mechanosensitive. The stiffness of the ECM surrounding blood vessels can vary significantly in space (location within the vascular tree), in time (aging) and in pathophysiological conditions (e.g. arteriosclerosis, cancer) (Chien et al., 2005; Collins et al., 2014; Krishnan et al., 2011; Volkman et al., 2002; Yeh et al., 2012). These variations in stiffness can dramatically affect gene expression and barrier integrity of VECs, thereby regulating the movement of vascular components from the bloodstream into underlying tissues, including the dissemination of bacterial pathogens (Lemichez et al., 2010). Clinical studies have shown that the local compliance of the basement membrane of VECs in the brain tissue is as soft as 1 kPa, while bigger vessels such as the aorta are as stiff as 20 kPa (Janmey and Miller, 2011; Kohn et al., 2015; Onken et al., 2014; Wells, 2008; Wood et al., 2010), and their stiffness can increase up to 70 kPa due to aging (Huynh et al., 2011) and in the context of cardiovascular diseases (CVDs), such as atherosclerosis or hypertension (Blacher et al., 1999; Boutouyrie et al., 2002). VECs, like other adherent cell types, probe features of their environment including its stiffness through integrins, transmembrane receptors that allow attachment of the cells to their matrix through direct binding to ECM ligand proteins (Schwartz, 2010; Senger et al., 2002). Integrin binding to the ECM leads to recruitment of additional proteins and to formation of focal adhesions that in turn relay information to the actin cytoskeleton and to various signaling molecules, modulating cellular adhesion, shape, contractility, gene expression and fate in general (Pelham and Wang, 1997). Focal adhesion kinase (FAK) is a non-receptor tyrosine kinase that has been established as a key component of the signal transduction pathways triggered by integrins. The expression and/or activity of FAK is dependent on ECM rigidity, showing decreased expression and/or activation on softer matrices for certain cell types and can influence cell adhesion and motility (Du et al., 2016; Higuita-Castro et al., 2014; Khatiwala et al., 2006; Provenzano et al., 2009). However, no studies have previously addressed if and how VEC matrix stiffness sensing through focal adhesions might affect host cells’ interactions with bacterial pathogens.

Bacterial adhesion to the surface of cells is typically the initial event in the pathogenesis of infection and can occur through receptor-mediated interactions between the host cell and the pathogen (Pizarro-Cerdá and Cossart, 2006). Vimentin is generally an intracellular cytoskeletal protein that forms intermediate filaments in many mesoderm-derived cells (Clarke and Allan, 2002). Vimentin can also be localized on the surface of cells for a variety of cell types, although the precise mechanism whereby vimentin is delivered to the surface of cells is not yet fully understood (Mitra et al., 2015; Mor-Vaknin et al., 2003; Päll et al., 2011; Rohrbeck et al., 2014; Shigyo et al., 2015). Recent studies have shown that various bacteria as well as viruses use surface vimentin as attachment receptor to facilitate entry into host cells (Bhattacharya et al., 2009; Du et al., 2014; Garg et al., 2006; Mak and Brüggemann, 2016; Rohrbeck et al., 2014; Yang et al., 2016; Yu et al., 2016; Zou et al., 2006). However, little is known about what modulates the amount of vimentin exposed on the surface of cells and whether regulation of its surface presentation potentially mediates bacterial uptake.

To assess whether ECM stiffness could mediate infection susceptibility of host cells, we used Lm as our model pathogen and examined *in vitro* Lm infection in endothelial cells derived from the microvasculature (human microvascular endothelial cells, HMEC-1) cultured on matrices of varying stiffness. Subendothelial stiffness depends on the blood vessel’s location within the body, on the size of vessel, on age and on physiological or pathophysiological conditions and so we chose our substrates’ stiffness to span over a large range of stiffnesses, from 0.6 kPa to 70 kPa (Kohn et al., 2015; Stroka and Aranda-Espinoza, 2011). We found a two-fold increase in bacterial adhesion when cells reside on stiff as compared to soft matrices. We also found that Tyr397 phosphorylation of FAK was higher for VECs residing on stiff matrices, and furthermore that knockdown of FAK or treatment of VECs with FAK inhibitors led to a decrease in bacterial infection. When we searched for candidate VEC surface receptors differentially modulated depending on FAK activity, we identified surface vimentin as a candidate, and found that decreasing the amount of host surface vimentin leads to a concomitant decrease in Lm infection. Taken together, our findings provide evidence that environmental stiffness of VECs, a previously unappreciated factor, plays an important role in mediating infection susceptibility to Lm through differential activity of FAK that in turn affects the amount of surface vimentin.

## RESULTS

### Uptake of *Listeria monocytogenes* by HMEC-1 is more efficient when cells reside on stiff substrates

Substrates on which vascular endothelial cells (VECs) are cultured *in vitro,* commonly glass or tissue culture (TC) polystyrene, are approximately six orders of magnitude stiffer than the natural ECM of human VECs (Dussurget et al., 2004; Sperling and Friedman, 1969). To be able to recapitulate *in vivo* conditions and determine in a systematic way the effect of the VECs’ matrix stiffness on the efficiency of *Listeria monocytogenes* (Lm) infection, we developed an assay based on manufacturing thin polyacrylamide hydrogels of varying stiffness ranging from 0.6 to 70 kPa on multi-well glass-bottom plates (Ahmed, 2015; Georges et al., 2006; Mih et al., 2011) (Suppl. Fig. 1A). To enable cell adhesion, hydrogels were surface-coated with collagen I, using a consistent density of collagen independent of hydrogel stiffness, as reported in previous studies (Huang et al., 2013; Tse and Engler, 2010; Yeung et al., 2005). Human microvascular endothelial cells (HMEC-1) seeded for 24 h on the hydrogels formed monolayers with comparable densities regardless of substrate stiffness, similar to growth of these cells on TC polystyrene (Suppl. Fig. 1B)

In order to measure the efficiency of bacterial infection quantitatively, we infected HMEC-1 with a strain of Lm lacking expression of the ActA protein, which is necessary for intra- and inter-cellular spread of this bacterium. In that way, we could attribute changes in the number of infected cells directly to initial invasion events and not to spread of the bacteria from cell to cell. We also confirmed that adhesion and invasion of wild-type Lm into HMEC-1 is similar to that of *ΔactA* Lm, consistent with previous studies on other host cell types (Brundage et al., 1993; Kocks et al., 1992), suggesting that ActA is not involved in adhesion or invasion (Suppl. Fig. 1 D-F). The *ΔactA* Lm strain we used also expresses a fluorescent protein under a promoter that is activated several hours after exposure of the bacteria to the host cell cytosol (actAp::mTagRFP) (Zeldovich et al., 2011), allowing reliable fluorescence-based detection only of bacteria that have successfully invaded the host cells. We determined the fraction of HMEC-1 infected with Lm in each well using flow cytometry (Fig. 1A-D). At a constant multiplicity of infection (MOI) and constant host cell density, we found a monotonic increase (strictly increasing relationship) in the number of VECs infected by Lm with increasing hydrogel stiffness that was highly reproducible among biological replicates (Fig. 1E and Suppl. Fig 1C). The overall efficiency of infection on the stiffest hydrogel substrates tested, 70 kPa, was comparable to the efficiency of infection for the same cells grown on TC polystyrene, suggesting that above a certain level of stiffness the infection efficiency does not change (Suppl. Fig 1C). Compared to this maximum level of infection for HMEC-1 cultured on the stiffest substrates, infection efficiency on the softest substrates decreased about two-fold (Fig. 1E and Suppl. Fig 1C).

**Figure 1.**
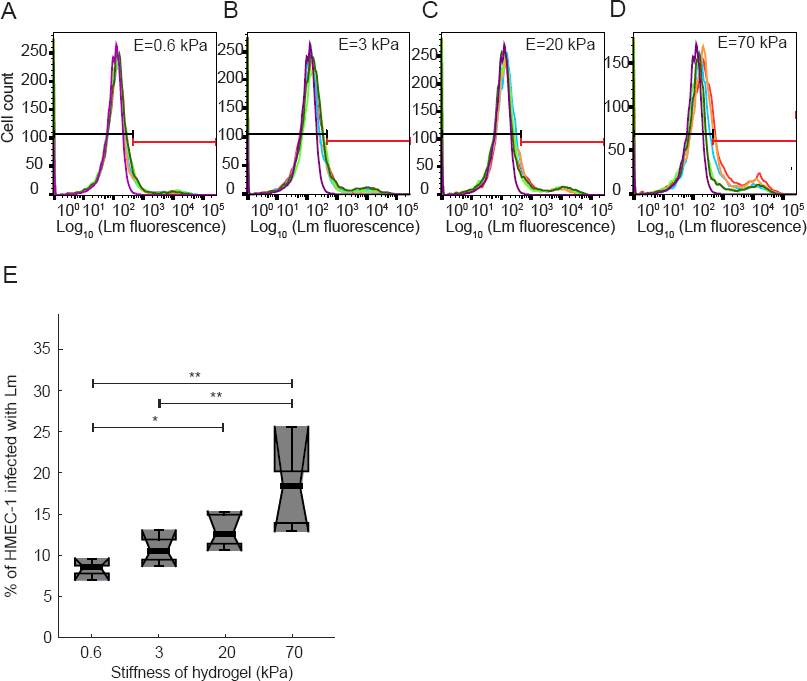
Uptake of Lm by HMEC-1 depends on the stiffness of the matrix on which cells reside. HMEC-1 residing on PA hydrogels of varying stiffness coated with collagen I were infected with *ΔactA* Lm (actAp::mTagRFP). Infection was analyzed by flow cytometry 7-8 h post infection. Bacteria were added at a multiplicity of infection (MOI) between 30 to 50 bacteria per host cell. (**A-D**) Histograms of the logarithm of bacterial fluorescence intensity per cell for HMEC-1 plated on 0.6 kPa (**A**), 3 kPa (**B**), 20 kPa (**C**) and 70 kPa (**D**) PA hydrogels. Histograms for N=5 replicates are shown in different colors. Histogram of control uninfected cells is shown in purple. Based on the autofluorescence of the control group a gate is defined (see black and red lines) showing what is considered uninfected (left, black line) and infected (right, red line). (**E**)Boxplots of percentage of HMEC-1 infected with *AactA* Lm versus hydrogel stiffness for the data shown in panels A-D. Circles represent outliers, and the boxplots’ notched sections show the 95% confidence interval around the median (Wilcoxon-Mann-Whitney test, for details about boxplots see Materials and Methods). One or two asterisks denote statistically significant differences between the medians of two distributions (<0.05 or <0.01, respectively; Wilcoxon rank sum test).

### HMEC-1 on soft hydrogels show decreased FAK activity and decreased FAK activity leads to less efficient Lm infection

Focal adhesion kinase (FAK) has been established as a key component of the signal transduction pathways triggered by integrins binding to the ECM, and for some cell types FAK expression and/or phosphorylation on residue Y397 have been shown to depend on mechanical cues sensed by cells, such as shear stresses imposed by fluid flow and/or ECM rigidity (Du et al., 2016; Higuita-Castro et al., 2014; Khatiwala et al., 2006; Mathieu et al., 2010; Provenzano et al., 2009). We therefore sought to determine whether FAK expression or activity could be modulated depending on the substrate stiffness where HMEC-1 reside. To that end, HMEC-1 were grown on soft 3 kPa hydrogels, stiff 70 kPa hydrogels and TC polystyrene substrates treated with vehicle control or PF573228 FAK inhibitor. Cells were then harvested and total amounts of FAK as well as pY397-FAK were determined using Western blot analysis. We found that increasing the stiffness of the underlying substrate from 3 kPa to 70 kPa resulted in increasing levels of pY397-FAK, but the levels of total FAK remained largely unchanged (Fig.2A-C). In addition, the levels of pFAK-Y397 were similar for cells residing on stiff 70 kPa hydrogels as compared to TC polystyrene substrates (Fig. 2A-C). Treating cells residing on polystyrene with the specific FAK inhibitor PF573228 led to a significant decrease of pY397- FAK while total FAK levels remained unchanged as compared to controls (Fig. 2A-C). Our results suggest that FAK phosphorylation at Y397 is modulated by the stiffness of the substrate on which HMEC-1 reside and therefore by different levels of cell-ECM interaction-induced focal adhesion signaling.

**Figure 2.**
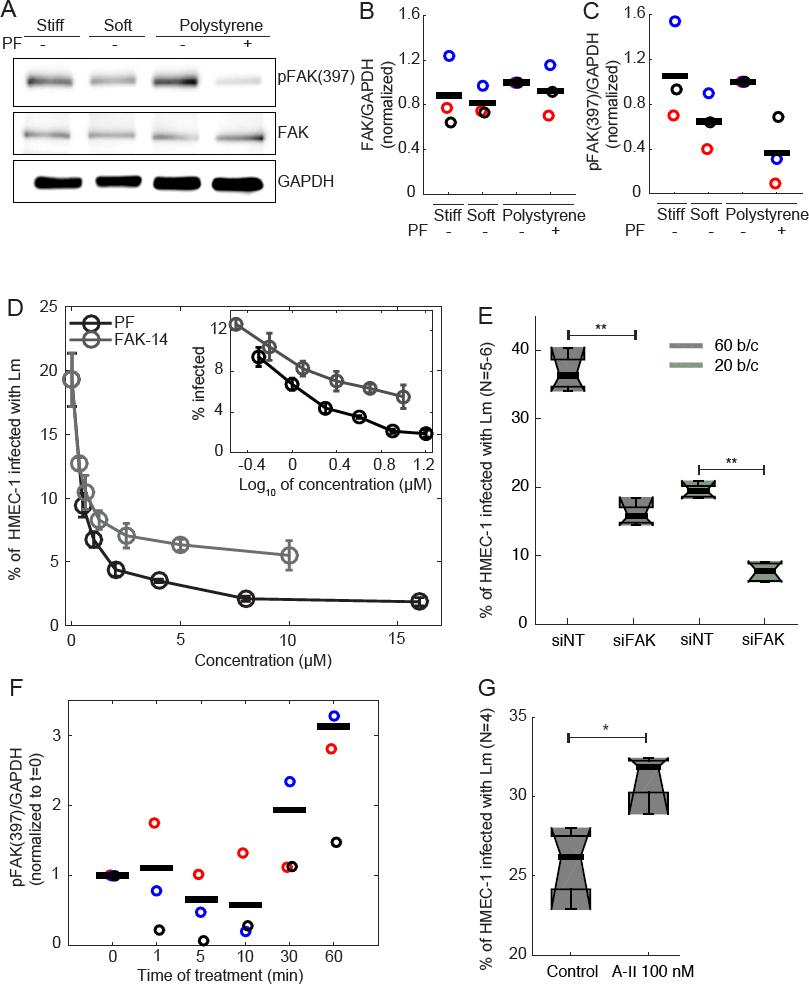
FAK activity of HMEC-1 residing on soft PA hydrogels is decreased, as is Lm uptake. (**A**) Western blots from whole HMEC-1 lysates showing expression of phosphorylated FAK (Tyr397) and total FAK for cells residing on soft gels (3 kPa), stiff gels (70 kPa) and TC polystyrene substrates with or without 2 μM PF537228 FAK inhibitor. In each Western blot, equal quantities of protein were loaded and equal loading was confirmed in relation to glyceraldehyde 3-phosphate dehydrogenase (GAPDH) expression. In each case, the Western blots shown are representative of 3 independent experiments. (**B-C**) Normalized ratio of FAK/GAPDH (**B**) and pFAK (Tyr397)/GAPDH (**C**) for HMEC-1 residing on varying stiffness substrates and treated or not with 2 μM PF537228 FAK inhibitor. Different color circles correspond to data from 3 independent experiments. Black bars represent the mean of the 3 independent experiments. For each experiment values have been normalized relative to the ratio for cells residing on polystyrene substrates. (**D**) Inhibition of bacterial uptake by FAK inhibitors. FAK-14, PF573228 or vehicle control was added 1 h before addition of bacteria to HMEC-1 residing on polystyrene substrates. Percentage of HMEC-1 infected with *ΔactA* Lm (actAp::mTagRFP) as a function of inhibitor concentration (mean +/− SD, N = 4 replicates). × = 0 corresponds to cells treated with vehicle control. Inset shows the same data with concentration on a log scale. Infection was analyzed by flow cytometry, 7-8 h after infection. MOI is 80. Representative data come from 1 of 3 independent experiments. (**E**) Boxplots of percentage of HMEC-1 infected with *ΔactA* Lm (actAp::mTagRFP) for cells treated either with non-targeting siRNA (siNT) or FAK siRNA (siFAK) (means +/−S.D., 3 independent experiments and N=6 replicates per experiment). MOI is 60 (gray) or 20 (green). Circles represent outliers, and the boxplots’ notched sections show the 95% confidence interval around the median (Wilcoxon-Mann-Whitney test, for details about boxplots see Materials and Methods). One or two asterisks denote statistically significant differences between the medians of two distributions (<0.05 or <0.01, respectively; Wilcoxon rank sum test). (**F**) Normalized ratio of pFAK (Tyr397)/GAPDH for HMEC-1 residing on polystyrene substrates and treated for varying amount of time (min) with 100 nM Angiotensin-II. Different color circles correspond to Western blot data from 3 independent experiments. In each Western blot, equal quantities of protein were loaded and equal loading was confirmed in relation to glyceraldehyde 3-phosphate dehydrogenase (GAPDH) expression. Black bars represent the mean of the 3 independent experiments. For each experiment values have been normalized relative to the ratio for untreated cells (t = 0 min). (**E**) Boxplots of percentage of HMEC-1 infected with *ΔactA* Lm (actAp::mTagRFP) for cells pre¬treated for 2 h either with vehicle control or 100 nM Angiotensin-II (means +/−S.D., 3 independent experiments and N=4 replicates per experiment). One or two asterisks denote statistically significant differences between the medians of two distributions (<0.05 or <0.01, respectively; Wilcoxon rank sum test).

To address whether reduced Lm uptake as observed for HMEC-1 residing on soft matrices can be attributed at least in part to reduced FAK activity, we treated HMEC-1 residing on polystyrene substrates with FAK inhibitors FAK-14 or PF573228 for one hour prior to infection. We then measured the efficiency of infection with Lm as described above, and found that both FAK inhibitors caused a similar dose-dependent inhibition of Lm infection (Fig. 2D). We then transfected HMEC-1 with commercial non-targeting control siRNA (siNT) or siRNA targeting FAK (siFAK) (Table S2) and total FAK expression was found to be about five-fold reduced for the FAK knockdown cells as compared to controls as determined by immunofluorescence (Suppl. Fig. 2A) and RT-PCR (Suppl Fig. 2B). We found a significant approximately 2-fold decrease in Lm infection efficiency for the FAK knockdown cells as compared to control cells (Fig. 2E; Suppl. Fig. 2C-D). Taken together these results are consistent with the hypothesis that decreased bacterial infection of HMEC-1 residing on highly compliant soft matrices may be in part due to decreased FAK activity.

To address whether elevating FAK activity can lead to increased bacterial uptake we pre-treated cells for varying times with vasoconstrictor peptide hormone Angiotensin II, which has been shown to induce an increase in phosphorylation of FAK *in vivo* in murine aortas (Louis et al., 2007) and also shown *in vitro* to lead to enhanced phosphorylation of FAK at Y397 for a variety of cell types (Greco et al., 2002; Torsoni et al., 2005; Weng and Shukla, 2002), including endothelial cells (Montiel et al., 2005). We first treated HMEC-1 with 100 nM Angiotensin II for varying times and then lysed the cells and used Western blotting to assess FAK expression and phosphorylation of FAK at Y397 (Fig. 2F and Suppl. Fig. 2F). Interestingly, we found that although FAK expression did not change upon treatment with Angiotensin II, phosphorylation of FAK at Y397 increased substantially for cells treated with Angiotensin II for more than 30 min. Since FAK activity was shown to be elevated for cells pretreated for more than 30 min with Angiotensin II, we then asked whether that would be sufficient to lead to increased infection susceptibility. When we infected HMEC-1 cells seeded on polystyrene substrates and pretreated with 100 nM of Angiotensin II, we observed a modest but significant 23% increase in infection susceptibility as compared to control cells (Fig. 2G and Suppl. Fig. 2E). We then seeded HMEC-1 cells on soft 3 kPa and stiff 70 kPa hydrogels and pretreated cells with or without 100 nM Angiotensin II for 2 h. We found a much stronger effect of Angiotensin II increasing infection for cells plated on soft substrates than on stiff substrates; indeed, treatment of cells on 3 kPa substrates with 100 nM Angiotensin II for 2 h was sufficient to raise the level of infection to be approximately equal to infection on 70 kPa substrates (Suppl. Fig. 2G). This result further supports the hypothesis that a primary determinant of the relatively inefficient infection of HMEC-1 on soft substrates is the relatively low level of FAK activity in this environmental condition.

### Adhesion but not invasion efficiency is increased when Lm infect HMEC-1 residing on stiff substrates

Decreased infection efficiency of Lm in HMEC-1 residing on soft matrices or treated with FAK inhibitors could be due to quantitative changes at several different steps in the infection process, for example decreased bacterial adhesion onto HMEC-1, or decreased invasion of the adhering bacteria into the HMEC-1, or a combination of both. To help identify the exact step in infection that is sensitive to substrate stiffness, we used a constitutively GFP-expressing strain of Lm to infect HMEC-1 residing on either soft 3 kPa hydrogels or stiff 70 kPa hydrogels, and treated with vehicle control or PF573228. Samples were fixed shortly after infection and bacteria that were attached to the cell surface but not yet internalized were specifically labeled with antibodies under non-permeabilizing conditions. This inside/outside labeling method allows us to distinguish between bacteria that are adhered but not internalized (GFP-positive and labeled with the antibody) and those that are fully internalized by the HMEC-1 (GFP-positive but not labeled with the antibody). We found that both the total number of bacteria per host cell and the number of internalized bacteria per cell are significantly increased when HMEC-1 reside on stiff 70 kPa hydrogels as compared to soft 3 kPa hydrogels (Fig. 3A-B). However, importantly, the invasion efficiency (ratio of internalized bacteria to total bacteria) did not differ between HMEC-1 residing on stiff as compared to soft matrices, suggesting that it is specifically the adhesion of bacteria to the surface of host cells that is increased when the matrix stiffness is elevated and which leads to increased Lm infection efficiency for cells residing on stiffer matrices (Fig. 3C). Interestingly, we found a decrease in both Lm adhesion and invasion efficiency irrespective of hydrogel stiffness for the cells treated with the FAK inhibitor (Fig. 3), suggesting that FAK activity levels modulate both adhesion and invasion efficiency of Lm in HMEC-1, while substrate stiffness affects adhesion only.

**Figure 3.**
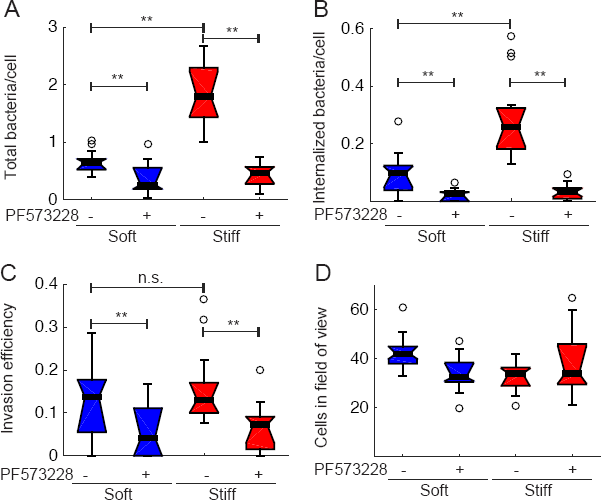
Lm adhesion but not invasion efficiency is increased when HMEC-1 reside on stiff hydrogels. HMEC-1 residing on soft (3 kPa) or stiff (70 kPa) PA hydrogels and treated with vehicle control or 2 μM PF537228 FAK inhibitor were infected with Lm (constitutively expressing GFP) at an MOI between 1.5 to 15. 30 min post-infection samples were fixed, and immunostained, and infection was analyzed by microscopy followed by image processing. Boxplots show: (**A**) total bacteria per cell; (**B**) internalized bacteria per cell; (**C**) invasion efficiency (ratio of internalized bacteria to total bacteria); (**D**) cells in the field of view. Representative data come from 1 of 3 independent experiments. N=800-1000 cells were analyzed for each condition. Two asterisks denote statistically significant differences between the medians of two distributions (<0.01; Wilcoxon rank sum test).

### InlB contributes to Lm infection of HMEC-1 but is not modulated by substrate stiffness

Our data indicate that lower FAK activity as it occurs when HMEC-1 are seeded on softer hydrogels leads to decreased Lm adhesion onto the surface of HMEC-1. This finding led us to hypothesize that a receptor at the surface of cells could be differentially regulated depending on substrate stiffness, leading to decreased bacterial adhesion. To examine whether any of the Lm internalins known to play a role in adhesion or invasion of other cell types, including VECs, is implicated in infection of HMEC-1 and responsible for the matrix stiffness dependent susceptibility to infection, we infected HMEC-1 with mutant strains deficient in each of the internalin-encoding genes *inlA, inlB* or *inlF* (Fig. 4A and Table S1). We found a two-fold decrease in bacterial infection when InlB was not present, suggesting that InlB is one of possibly multiple bacterial factors that contribute to Lm infection of HMEC-1. In contrast to the effect of the deletion in the *inlB* gene, deletion of *inlF* had no effect in bacterial infection of HMEC-1. Deletion of *inlA* resulted in very modest but reproducible reduction in infection as reported previously for human brain microvascular endothelial cells infected with Lm (Greiffenberg et al., 1998).

**Figure 4.**
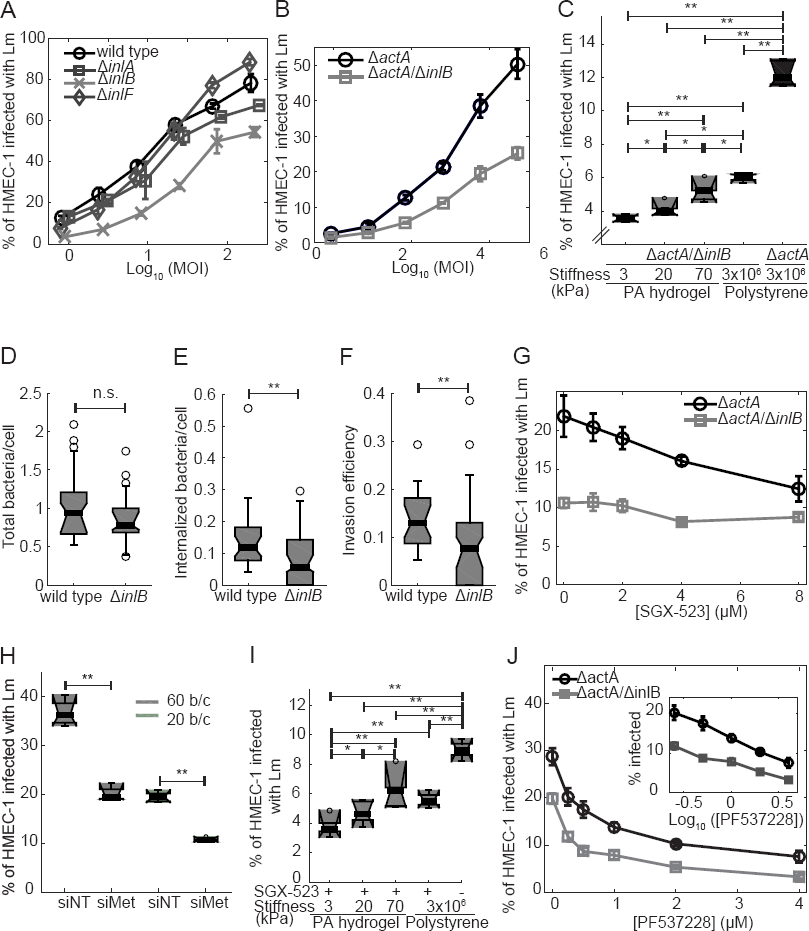
Infection of HMEC-1 by Lm is in part mediated by InlB in a manner independent of matrix stiffness. (A) Percentage of HMEC-1 infected with Lm as a function of the logarithm of MOI (mean +/− SD, N = 4 replicates). HMEC-1 were infected with the indicated strains: wild type (circle); *AinlA* (square); *AinlB* (cross); *ΔinlF* (diamond) (actAp::mTagRFP). The frequency of infected HMEC-1 was determined by flow cytometry 7-8 h post infection. Representative data come from 1 of 3 independent experiments. (B) Percentage of HMEC-1 infected with Lm as a function of the logarithm of MOI (mean +/− SD, N = 4 replicates). HMEC-1 were infected with the indicated strains: *AactA* (black circle); *ΔactA/ΔinlB* (gray square) (actAp::mTagRFP). The frequency of infected HMEC-1 was determined by flow cytometry 7-8 h post infection. Representative data come from 1 of 3 independent experiments. (C) Boxplots of percentage of HMEC-1 infected with Lm as a function of substrate stiffness (N = 5-6 replicates). HMEC-1 were infected with the indicated Lm strains: *AactA* (gray); *AactA/AinlB* (black) (actAp::mTagRFP) at an MOI of 20. Infection was analyzed by flow cytometry, 7-8 h after infection. Representative data come from 1 of 3 independent experiments. One or two asterisks denote statistically significant differences between the medians of two distributions (<0.05 or <0.01, respectively; Wilcoxon rank sum test). (**D-F**) HMEC-1 residing on collagen I-coated glass substrates were infected with Lm (constitutively expressing GFP) or *ΔinlB* Lm at an MOI of 3.5. 30 min post-infection samples were fixed, immunostained and infection was analyzed by microscopy followed by image processing. Boxplots showing: (**D**) total bacteria per cell; (**E**) internalized bacteria per cell; (**F**) invasion efficiency (ratio of internalized bacteria to total bacteria). For each condition, 500-550 cells were analyzed in total. (**G**) Percentage of HMEC-1 infected with Lm as a function of SGX-523 Met inhibitor concentration (mean +/− SD, N = 4 replicates). SGX-523 or vehicle control was added 1 h before addition of bacteria. HMEC-1 were infected with the indicated strains: *ΔactA* (black circle); *ΔactA/ΔinlB* (gray square) (actAp::mTagRFP) at an MOI of 75. Infection was analyzed by flow cytometry, 7-8 h after infection. Representative data come from 1 of 3 independent experiments. (**H**) Boxplots of percentage of HMEC-1 infected with *ΔactA* Lm (actAp::mTagRFP) for cells treated either with non-targeting siRNA (siNT) or Met siRNA (siFAK) (means +/−S.D., 3 independent experiments and N=6 replicates per experiment). MOI is 60 (gray) or 20 (green). Circles represent outliers, and the boxplots’ notched sections show the 95% confidence interval around the median (Wilcoxon-Mann-Whitney test, for details about boxplots see Materials and Methods). One or two asterisks denote statistically significant differences between the medians of two distributions (<0.05 or <0.01, respectively; Wilcoxon rank sum test). (**I**) Boxplots of percentage of HMEC-1 infected with Lm as a function of substrate stiffness (N = 5-6 replicates). HMEC-1 were treated with vehicle control or 1 μM SGX-523 Met inhibitor for 1 h prior to infection and then infected with *ΔactA* Lm (actAp::mTagRFP). Infection was analyzed by flow cytometry, 7-8 h after infection. MOI is 20. Representative data come from 1 of 3 independent experiments. (**J**) Percentage of HMEC-1 infected with Lm as a function of PF537228 inhibitor concentration (mean +/− SD, N = 4 replicates). PF537228 or vehicle control was added 1 h before addition of bacteria (see Suppl. Fig. 1D). HMEC-1 were infected with the indicated strains: *ΔactA* (black circle); *ΔactA/ΔinIB* (gray square) (actAp::mTagRFP) at an MOI of 75. Infection was analyzed by flow cytometry, 7-8 h after infection. Representative data come from 1 of 3 independent experiments. Inset shows the same data with concentration on a log scale.

The importance of InlB for initial invasion of Lm into HMEC-1 was further confirmed by infecting HMEC-1 with a different strain that had deletions in both *inlB* and *actA* so that bacteria could not spread from cell to cell. Consistent with our previous findings, we observed a two-fold decrease in bacterial uptake for a series of different multiplicities of infection (MOI) when *inlB* was knocked out (Fig. 4B). Importantly, bacteria carrying a deletion of *inlB* still exhibited a strong monotonic dependence of infection efficiency on substrate stiffness (Fig. 4C). This finding indicates that, while InlB contributes to some step in the infection of HMEC-1 by Lm, it is not governing the step that is sensitive to substrate stiffness.

We confirmed this conclusion using four different lines of experimentation. First, we directly assessed the effect of *inlB* deletion on bacterial adhesion and invasion of HMEC-1 using inside/outside labeling as described above. We found that the total number of bacteria associated with HMEC-1 cells was the same for wild-type bacteria as compared to an isogenic *ΔinlB* strain (Fig. 4D), that is, adhesion to the host cell surface is indeed unchanged. However, the number of internalized bacteria per cell and therefore the overall invasion efficiency was significantly decreased for the *ΔinlB* strain (Fig. 4E-F). Second, we examined the effect of inhibiting the host cell binding partner for InlB, the receptor tyrosine kinase Met (also known as the hepatocyte growth factor receptor), on bacterial invasion. We confirmed that HMEC-1 express Met (Suppl. Fig. 3A), and then treated the cells with varying concentrations of the highly selective Met inhibitor SGX-523 prior to infection (Buchanan et al., 2009). We found that only uptake of the control *ΔactA* strain of Lm decreased with increasing concentration of SGX-523, while uptake of the isogenic *ΔactA/ΔinlB* Lm did not change substantially as a function of SGX-523 concentration (Fig. 4G). Third, when we then transfected HMEC-1 with commercial non targeting control siRNA (siNT) or siRNA targeting Met (siMet) and then infected HMEC-1 with Lm, we found a significant 2-fold decrease in Lm infection efficiency for the Met knockdown cells as compared to control cells (Fig. 4H; Suppl. Fig. 3B-C), while confirming that Met expression was five-fold reduced for the Met knockdown cells as compared to controls through RT-PCR (Suppl. Fig. 3D and Table S2). When we then infected transfected siNT and siMet HMEC-1 with *ΔactA/ΔinlB* Lm, we found no significant difference in infection Lm uptake between the two conditions, consistent with the previous results of cells pretreated or not with the SGX-523 (Fig. 4G and Suppl. Fig. 3E). These results confirm that InlB most likely contributes to Lm invasion into HMEC-1 by interacting with host Met, as has been shown for other cell types (Bierne and Cossart, 2002; Parida et al., 1998). However, Lm invasion into HMEC-1 treated with SGX-523 still retains a strong dependence on substrate stiffness (Fig. 4I), just as we found for the *ΔinlB* strain. Fourth, we found that infection of HMEC-1 with the *ΔactA/ΔinlB* strain of Lm is still highly sensitive to inhibition of FAK with PF573228, showing a dose-response curve that parallels that of the control *ΔactA* strain (Fig. 4J).

So far we can conclude that substrate stiffness affects the adhesion of Lm to the surface of HMEC-1 such that bacterial adhesion to host cells residing on soft substrates is about two fold less than to host cells residing on stiff substrates. The well-characterized InlB-Met interaction appears to contribute to invasion of adherent bacteria into HMEC-1, and is responsible for about half of the invasion efficiency for Lm infecting these cells, however this interaction does not govern bacterial adhesion per se, is not sensitive to substrate stiffness, and is orthogonal to the FAK signaling axis. We therefore sought to identify alternative possible host cell receptors for adhesion of Lm to the surface of HMEC-1 that might be responsible for the stiffness-dependent effects we consistently observe.

### Inhibiting FAK activity leads to a reduced amount of cell surface vimentin for HMEC-1

Since the decreased FAK activity associated with growth of HMEC-1 on softer hydrogels leads to decreased Lm adhesion onto the surface of host cells, we attempted to identify candidate host surface receptors whose amount is modulated by FAK activity and which mediate bacterial adhesion. Since manufacturing soft and stiff PA hydrogels with large dimensions is both technically challenging and costly, we instead grew cells on large TC polystyrene plates and treated them with vehicle control or the FAK inhibitor PF573228. We then biotinylated the cell surface-exposed proteins using membrane-impermeable EZ-Link Sulfo-NHS-SS-Biotin and isolated them by affinity chromatography with streptavidin-agarose beads (Fujimoto et al., 1992; Nunomura et al., 2005). Isolated surface proteins were then run in 2D-PAGE gels to allow separation of proteins according to both their isoelectric points and their molecular weight (Elia, 2012). Gels were silver stained and comparison between gels was performed by marking the spots that differed in intensity (Fig. 5A-B). We found a spot at approximately 55 kDa consistently differing in two independent experiments between the control sample and the sample treated with the FAK inhibitor PF573228 (see circles in Fig. 5A-B). In a third replicate experiment of surface protein isolation, 2D-PAGE gels were Coomassie stained and that specific 55 kDa spot that differed as a function of FAK inhibitor treatment was excised and further analyzed by nanoliquid chromatography tandem mass spectrometry (nanoLC-MS/MS). We identified this spot as the protein vimentin (accession number: gi|340219, Mascot identification score: 788, number of identified peptides: 87, mass: 53752, isoelectric point: 5.05, sequence coverage: 61%). Vimentin is a major component of the cytoskeleton forming intermediate filaments, but exists also in soluble form and has been previously found to localize at the cell surface for a wide variety of cells including VECs (Du et al., 2014; Fuchs and Weber, 1994; Mitra et al., 2015; Yu et al., 2016; Zou et al., 2006).

**Figure 5.**
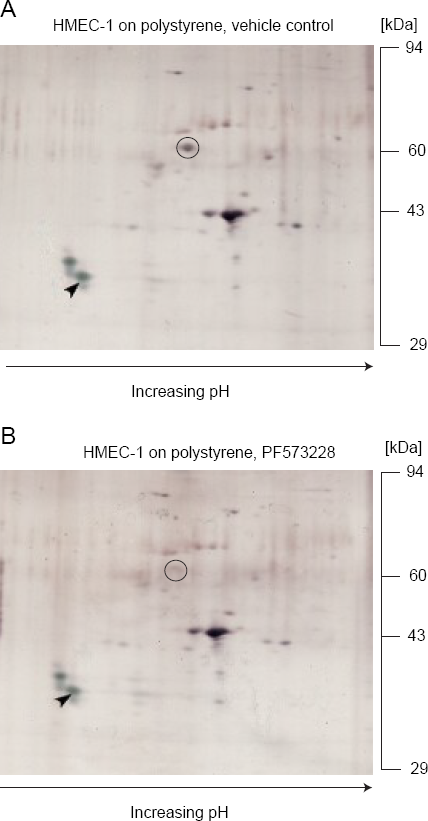
Lower FAK activity leads to reduced amount of cell surface vimentin. (**A-B**) 2D-PAGE gels of plasma membrane proteins of HMEC-1 grown on TC polystyrene substrates treated for 1 h with vehicle control (**A**) or 2 μM PF537228 FAK inhibitor (**B**). pH increases from left to right. Gels were silver stained and one isoelectric point marker (tropomyosin), added to each sample as an internal standard, is marked with a black arrow. The one spot that differed consistently between 3 independent experiments is indicated with a black circle and corresponds to vimentin (55 kDa).

Given the high validity of identification, we sought to confirm whether vimentin is expressed at the surface of HMEC-1 by performing immunostaining of non-permeabilized HMEC-1 seeded on collagen I-coated glass substrates followed by epifluorescence imaging (Fig. 6). For negative controls, cells were incubated with secondary antibody alone. The anti-vimentin H-84 antibody stained the vimentin intermediate filaments of permeabilized cells very well (Fig. 6A-B). Non-permeabilized cells showed surface staining of vimentin near cell-cell junctions and very low background staining was observed for cells incubated with secondary antibody alone (Fig. 6C-D). This localization confirmed that cell surface vimentin could be a plausible candidate responsible for adhesion of Lm to HMEC-1.

**Figure 6.**
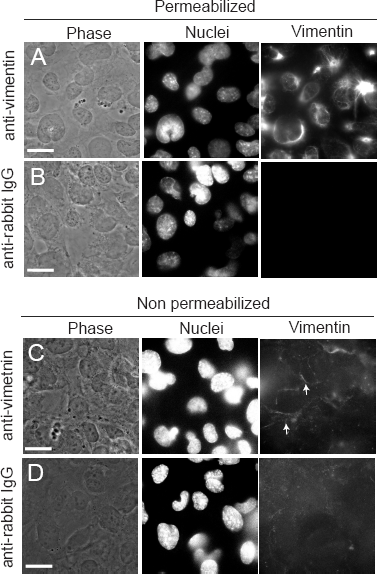
Surface vimentin is localized along the periphery of HMEC-1. (**A-D**) Cells were stained for vimentin using the rabbit anti-vimentin H-84 antibody. For negative controls, cells were stained with secondary anti-rabbit IgG antibody alone. Representative phase image of cells (left column), image of the nuclei (middle column) and H-84 anti-vimentin antibody fluorescence (right column) are shown for: (**A**) Permeabilized HMEC-1 strained for intracellular vimentin; (**B**) Permeabilized HMEC-1 incubated with anti-rabbit IgG alone as negative controls. (**C**) Non permeabilized HMEC-1 stained for surface vimentin; (D) Non permeabilized HMEC-1 incubated with anti-rabbit IgG alone as negative controls. Scale bar shown in white is 20 μm. White arrows point at the localization of surface vimentin at cell-cell junctions.

### HMEC-1 surface vimentin contributes to Lm uptake

To determine whether surface vimentin specifically contributes to Lm adhesion and thereby infection of HMEC-1, we used three independent methods to decrease the availability of cell surface vimentin for bacterial adhesion. First, we pretreated HMEC-1 with varying concentrations of H-84 anti-vimentin antibody prior to infection with Lm, and found a dose-dependent decrease in infection efficiency (Fig. 7A). We found a maximum two-fold decrease in Lm infection when incubating cells with 80 μg/ml anti-vimentin antibody, while increasing further the concentration of the antibody did not lead to further decrease in Lm uptake (Suppl. Fig. 3A).

**Figure 7.**
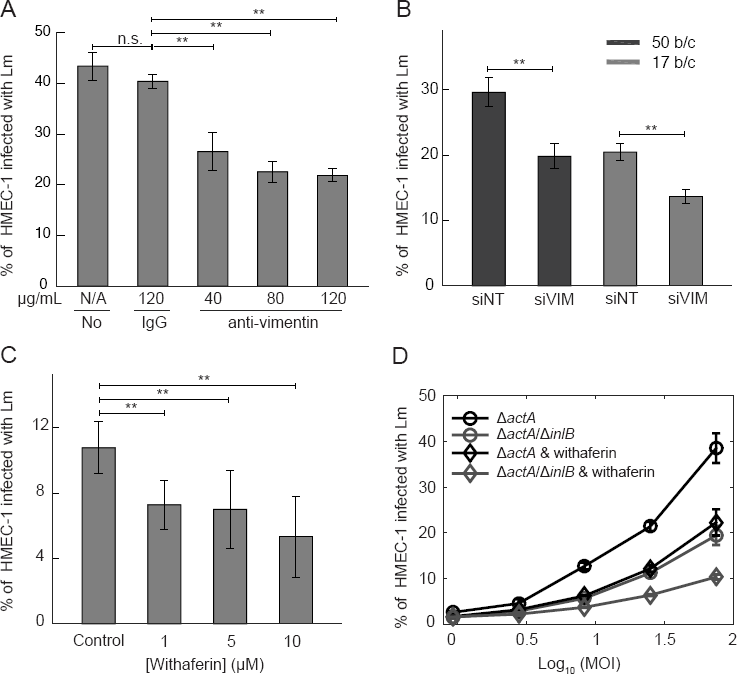
Surface vimentin of HMEC-1 is implicated in Lm uptake. (**A**) Decrease in bacterial uptake after blocking HMEC-1 with anti-vimentin antibody H-84. Barplots of percentage of HMEC-1 infected with *ΔactA* Lm (actAp::mTagRFP) as a function of antibody concentration (means +/−S.D. and N=6 replicates per experiment). Representative data come from 1 of 3 independent experiments. Infection was analyzed by flow cytometry, 7-8 h after infection. (**B**) Barplots of percentage of HMEC-1 infected with *ΔactA* Lm (actAp::mTagRFP) for cells treated either with non-targeting siRNA (siNT) or vimentin siRNA (siVIM) (means +/−S.D., and N=6 replicates per experiment). Representative data come from 1 of 3 independent experiments.MOI is 50 (black barplots) and 17 (gray barplots). (**C**) Decreased uptake of Lm when HMEC-1 are treated with withaferin that captures soluble vimentin 30 min prior to infection. Barplots of percentage of HMEC-1 infected with *ΔactA* Lm (actAp::mTagRFP) as a function of withaferin concentration (means +/−S.D. and N=6 replicates per experiment). Representative data come from 1 of 3 independent experiments. Infection was analyzed by flow cytometry, 7-8 h after infection. (**D**) Percentage of HMEC-1 infected with Lm as a function of the logarithm of MOI (mean +/− SD, N = 4 replicates). HMEC-1 were infected with the indicated strains: *ΔactA* (black); *ΔactA/ΔinlB* (gray) (actAp::mTagRFP) and HMEC-1 were treated with vehicle control (circle) or withaferin (diamond) for 30 min prior to infection. The frequency of infected HMEC-1 was determined by flow cytometry 7-8 h post infection. Representative data come from 1 of 3 independent experiments. MOI ranged from 50 to 120. One or two asterisks denote statistically significant differences between the medians of infection fraction of control versus all other groups (<0.05 or <0.01, respectively; Wilcoxon rank sum test).

Second, HMEC-1 were transfected with commercial non-targeting control siRNA (siNT) or siRNA targeting vimentin (siVIM) and total vimentin expression was found to be five-fold reduced for the vimentin knockdown cells as compared to controls through RT-PCR and reduced total vimentin expression was also confirmed through immunofluorescence (Suppl. Fig. 4A-B and Table S2). We found a significant decrease (about 30%) in Lm infection efficiency for the vimentin knockdown cells as compared to control cells, while confirming that cell confluency under both conditions was similar (Fig. 7B; Suppl. Fig. 4C-F).

Third, we treated HMEC-1 cells with varying concentrations of withaferin, a natural product known to bind the soluble form of vimentin (Bargagna-Mohan et al., 2013; Bargagna-Mohan et al.; Chi et al., 2010). We found a dose-dependent decrease in infection efficiency for Lm in withaferin-treated cells, again finding a maximum two-fold decrease when cells were treated with 5-10 μM of withaferin prior to infection (Fig. 7C). Higher concentrations of withaferin disrupted HMEC-1 cell-substrate adhesion. Overall these results are consistent with the hypothesis that cell surface vimentin is responsible for about half of the amount of adhesion of Lm to the HMEC-1 surface, and that vimentin presentation on the cell surface is sensitive to substrate stiffness.

As an additional control to confirm that anti-vimentin antibodies and withaferin treatment inhibit Lm uptake due to effects on surface vimentin and not to non-specific effects, we transfected HMEC-1 with commercial non-targeting control siRNA (siNT) or siRNA targeting vimentin (siVIM) and then blocked cells for 1h with the H-84 anti-vimentin antibody. We then infected the host cells with constitutively GFP-expressing Lm and fixed samples shortly after infection. Through quantitative microscopy we observed no significant change in infection susceptibility for vimentin knock-down cells blocked with anti-vimentin antibodies prior to infection as compared to vimentin knock-downs alone, confirming that the antibodies inhibit uptake due to effects on vimentin and not to non-specific effects (Suppl Fig. 4G). Similarly, when HMEC-1 transfected with siNT or siVIM were pretreated with our without withaferin we found a significant decrease in bacterial adhesion only for the control cells treated with withaferin and not for the vimentin knock-down cells (Suppl Fig. 4H). Finally, we hypothesized that if vimentin localizes at cell-cell junctions, and given that perturbing the amount of surface vimentin leads to up to a 2-fold decrease in infection susceptibility, then we should expect that when infecting HMEC-1 with Lm half of the bacteria should adhere at cell-cell junctions and half of the bacteria could adhere anywhere else. Indeed, when we infected HMEC-1 with Lm, fixed the samples and then immunostained for VE-cadherin as a proxy for cell-cell junctions (Suppl. Fig. 5A), we observed that quite often bacteria were found adhering at cell-cell junctions. This was not the case for vimentin knockdown cells where bacteria where found all across the apical surface of cells (Suppl. Fig. 5B).

To examine any possible interaction between the contribution of vimentin-based adhesion and the contribution of InlB/Met-based invasion of Lm infecting HMEC-1, we pretreated HMEC-1 with withaferin or vehicle control and then infected with either *ΔactA* or *ΔactA/ΔinlB* Lm over a range of multiplicities of infection (MOI). Consistent with our results described above, we found a two-fold decrease in Lm uptake when cells were pretreated with withaferin and infected with *ΔactA* Lm or when cells were not treated with withaferin but were infected with *ΔactA/ΔinlB* Lm (Fig. 7D). When cells were both pretreated with withaferin and infected with *ΔactA/ΔinlB* Lm, infection was decreased four-fold relative to control (Fig. 7D), i.e. InlB/Met and vimentin contribute independently to infection efficiency. However, under all conditions there was still residual infection, suggesting that Lm uses additional strategies to achieve uptake by HMEC-1, other than interactions with surface vimentin for bacterial adhesion and with Met for increased bacterial internalization.

### Uptake of *Listeria innocua* but not carboxylated latex beads by HMEC-1 depends on cell surface vimentin

Our results demonstrate that the vimentin-dependent adhesion of Lm to HMEC-1 is sensitive to substrate stiffness and is not mediated by the bacterial internalins InlA, InlB, or InlF. In order to gain insight into whether vimentin-based bacterial adhesion is a result of a non specific stickiness or a specific host-bacterium interaction, we measured the influence of cell surface vimentin on adhesion of other bacteria and of non-biological particles. *Listeria innocua* (Li) is a bacterium closely related to Lm that is considered non-pathogenic in that it lacks the putative internalin family members and other virulence factors that Lm carries (Glaser et al., 2001; Lauer et al., 2002). When we infected HMEC-1 with comparable loads of Lm and Li, we found that adhesion of both bacteria is comparable (Fig. 8A). We also found that treating HMEC-1 with FAK inhibitor PF537228 or blocking HMEC-1 with anti-vimentin antibody H-84 reduces adhesion of both Lm and Li onto HMEC-1 by similar amounts (Fig. 8A). Next, to examine whether a bacterial factor common in Lm and Li could responsible for surface vimentin mediated adhesion, we blocked HMEC-1 with anti-vimentin antibody and exposed cells to 2 μm carboxylated latex beads, which are efficiently taken up by VECs. We found that uptake of beads did not vary with antibody concentration and was identical to uptake of cells treated with isotype control (Fig. 8B). Our results suggest that cell surface vimentin is a common receptor for both Lm and Li in the context of adhesion to HMEC-1, but uptake of nonbiological particles such as carboxylated latex beads does not depend on surface vimentin.

**Figure 8.**
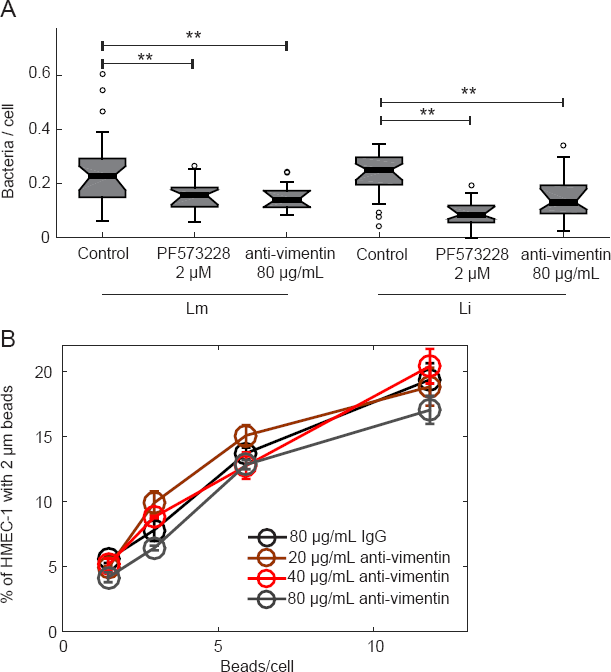
Blocking HMEC-1 with anti-vimentin antibody reduces Li adhesion onto HMEC-1xg375. (**A**) Boxplots showing the number of bacteria per cell, for HMEC-1 residing on glass substrates and treated with vehicle control, 2 μM PF537228 FAK inhibitor, or 80 μg/mL H-84 anti-vimentin antibody prior to infection. Cells were infected with Lm or Li at an MOI of 4. 30 min post-infection samples were fixed, immunostained and adhesion of bacteria was analyzed by microscopy followed by image processing. For each condition, 2300-2600 cells were analyzed in total and data refer to 1 of 2 independent experiments. Two asterisks denote statistically significant differences between the median values of control cells versus all other groups <0.01; Wilcoxon rank sum test). (**B**) HMEC-1 residing on TC polystyrene substrates and blocked for 1 h with varying concentrations of H-84 anti-vimentin antibody or isotype control were “infected” with 2 μm beads at different concentrations. The frequency of micro-bead uptake by HMEC-1 was determined by flow cytometry 2 h post addition of beads. Plot shows percentage of cells that internalized beads as a function of the beads/cell added for different H-84 antibody concentrations (mean +/− SD, N = 4 replicates).

## DISCUSSION

### Bacterial infection of VECs depends on the stiffness of the matrix on which cells reside

Our study provides the first evidence that ECM stiffness is a crucial determinant in modulating susceptibility of host cells to infection by bacterial pathogens. Using HMEC-1 cells as model host adherent cells and Lm as model bacterial pathogen, we demonstrated that host cell susceptibility to Lm infection increases with increasing ECM stiffness. Mechanosensitive cells, such as VECs, can “feel” a variety of physical cues such as fluid shear stresses, cyclic mechanical strains, matrix stiffness and respond to them by altering their morphology, gene expression and function (Kohn et al., 2015; Reinhart-King, 2008). Interestingly, many signaling pathways involved in mechanotransduction are common irrespective of the exact mechanical stimulus (Li et al., 1997; Zebda et al., 2012). For instance, exposure of VECs to laminar shear stress leads to recruitment of FAK at focal adhesions and to an increase in the phosphorylation at Y397 (Li et al., 1997). Consistent with FAK activity being sensitive to mechanical stimuli, we found that FAK activity is increased when cells reside on stiffer matrices. Concurrently, we observe a decrease in Lm uptake when FAK activity is reduced or inhibited. It is possible that enhanced FAK activity for VECs exposed to shear stresses, might also lead to increased Lm uptake, but this still remains to be investigated (Li et al., 1997). In our study, we only focused on the effect of ECM stiffness on host cell infection susceptibility and did not consider in our model additional mechanical cues that host cells experience (Shi and Tarbell, 2011). Based on our findings and on recent studies on shear flow-exposed endothelial cells infected with pathogens *B. burgdorferi* or S. *aureus,* we speculate that similar to matrix stiffness it is plausible that additional mechanical cues could also play an important role in mediating susceptibility of adherent host cells to bacterial infection (Claes et al., 2014; Niddam et al., 2017). Therefore, the development of organotypic models for studying bacterial infection, that take into account all of the different *in vivo* mechanical cues sensed by cells, is pertinent and will aid our emerging understanding of the biomechanics of host-bacteria interactions as well as the design of more effective therapeutic interventions that take into consideration the local cellular mechanical micro-environment.

Our findings suggest that together with biochemical cues, the stiffness of the environment where cells reside is an additional variable worth considering since it affects cellular behavior and susceptibility to bacterial infection. In traditional studies of bacterial infection of host cells’ ECM stiffness is not taken into account and such studies are commonly performed on non-physiological glass or polystyrene surfaces. Our study on how HMEC-1 interact with Lm when varying subendothelial stiffness was designed with the intent to mimic variations that occur in physiological conditions depending on the precise anatomic location of the vessel, in aging and during pathophysiological conditions such as cardiovascular disease (Zieman et al., 2005) or cancer (Acerbi et al., 2015). Our lower range of subendothelial matrix stiffness (0.6 kPa) could be relevant for the brain microvasculature where the blood vessel microenvironment is very soft whereas stiffer values are more relevant for bigger vessels or aged/diseased vasculature (Kohn et al., 2015; Stroka and Aranda-Espinoza, 2011). For instance, clinical studies on arteries of atherosclerotic mice have shown that vessel stiffness for diseased mice can increase from 5 kPa to up to 28 kPa (Kothapalli et al., 2012). Similarly, injury can increase vessel stiffness as has been shown for injured femoral artery whose stiffness increases from 3 kPa to 10 kPa (Klein et al., 2009). Finally, studies have shown that aging can lead to significant vessel stiffening with studies suggesting that the aorta stiffness due to aging can raise from 20 kPa to 70 kPa (Huynh et al., 2011). We showed that subendothelial stiffness changes the amount of surface vimentin expressed by VECs, therefore affecting critically how much host cells will be susceptible to bacterial infection. That is atherosclerotic vessels or aged vessels might be more susceptible to infection based on our findings, while VECs lining healthy and softer vessels should be less susceptible to infection by Lm. It is plausible that low degree of stiffness might be a host mechanism to protect against infection. For the purpose of the present study we just examined how subendothelial stiffness modulates bacterial adhesion on previously uninfected VECs. However, it is very possible that once bacteria infect their hosts they might have developed ways to circumvent such a host mechanism by somehow altering matrix stiffness, *i.e.* by reprogramming host cells to alter their production of ECM proteins or matrix-degrading enzymes, which could lead to alterations in subendothelial stiffness and subsequently VEC biomechanics and contractility, critically impacting infection. Future studies should address how host cell biomechanics change upon infection and how bacterial infection changes susceptibility of host cells to further infection.

### Surface vimentin is a receptor for *Listeria* infection of VECs

We found that lower FAK activity as occurs when HMEC-1 are seeded on softer hydrogels leads to decreased Lm and Li adhesion onto the surface of HMEC-1 and that is attributable to lower levels of cell surface vimentin. The role of cell surface vimentin as an attachment receptor facilitating bacterial or viral entry has been previously documented for other pathogens (Bhattacharya et al., 2009; Du et al., 2014; Garg et al., 2006; Mak and Brüggemann, 2016; Rohrbeck et al., 2014; Yang et al., 2016; Yu et al., 2016; Zou et al., 2006). VEC surface vimentin in particular has been shown to be the receptor for attachment of *E. coli* K1 and multiple viruses (Das et al., 2011; Du et al., 2014; Koudelka et al., 2009; Yang et al., 2016; Zou et al., 2006). We speculate that matrix stiffness of VECs might possibly also affect uptake of the above viruses and bacteria, since the amount of surface vimentin is reduced when FAK activity is decreased as in the case of cells residing on highly compliant matrices.

Bacterial adhesion to the surface of cells is commonly required as a precursor to invasion (Lebrun et al., 1996; Lecuit et al., 2000). Our finding that both Lm and Li adhesion is reduced when HMEC-1 are blocked with anti-vimentin antibodies suggests that the possible bacterial binding partner of vimentin might be expressed in both bacterial species. Surface vimentin has been shown to possess lectin-like activity binding N-acetylglucosamine (GlcNAc), a sugar that is also commonly found as a component of bacterial cell envelopes (Ise et al., 2010; Konopka, 2012). Specifically, GlcNAc is found decorating the wall teichoic acids of Lm serotype 1/2a strains, such as the 10403S strain used in this study (Fiedler, 1988). That implies that bacterial carbohydrate-binding to host cell surface vimentin might also be a plausible specific interaction.

### Lm uses multiple pathogenic strategies to infect VECs, some of which are independent of ECM stiffness

We identified that the Lm virulence factor InlB contributes to invasion of HMEC-1 cells, mostly likely via its well-characterized interaction with the host cell receptor Met. Similarly, Lm uptake is decreased by two-fold when Met is inhibited. However deletion of *inlB* and decrease of Met activity by siRNA knockdown or by pharmacological inhibition do not affect the ability of the infection process to respond to changes in substrate stiffness. When we infected HMEC-1 with *ΔactA/ΔinlB* Lm after treating host cells with withaferin that binds to soluble vimentin, we found a four-fold decrease in infection relative to controls, indicating that the vimentin-dependent and InlB-dependent pathways contribute to infection independently of one another; however, even when both pathways were blocked there was still residual Lm infection. This residual Lm infection suggests that there are additional InlB-independent and vimentin-independent mechanisms that Lm leverage to infect HMEC-1.

Increased Lm adhesion onto HMEC-1 residing on stiff matrices could be attributed to multiple factors. In this work, we investigated the possibility of a receptor at the surface of host cells being differentially regulated when FAK activity is increased, as in stiff matrices, and identified cell surface vimentin as a well-supported candidate receptor. However, it is also possible that additional factors might also lead to the decreased adhesion of Lm onto HMEC-1 residing on softer matrices. It is plausible that the cell surface glycans of HMEC-1 might be differentially regulated depending on the stiffness of the environment. For example previous studies have suggested both a positive correlation between FAK activity and the heparan sulfate portion of the glycocalyx (Gopal et al., 2010; Liu et al., 2002) and between the heparan sulfate portion of the glycocalyx and Lm adhesion onto enterocytes (Henry-Stanley et al., 2003). Another possibility could be the differential regulation of biophysical or mechanical properties of the host cells depending on matrix stiffness (e.g. cell cortical stiffness, surface roughness, membrane tension). For instance, studies on adhesion of bacteria onto hydrogel surfaces have shown that adherence is increased on stiffer hydrogels (Kolewe et al., 2015), suggesting that if cells become stiffer on stiffer hydrogels, cellular stiffness alone could lead to increased bacterial adhesion (Higuita-Castro et al., 2014; Tang et al., 2012).

### VEC origin and physical cues might explain discrepancy between previous studies on VEC infection by Lm

The endothelial lining of blood vessels displays remarkable spatiotemporal heterogeneity and the basis of these morphological and functional differences are still unclear (Aird, 2007; Aird, 2012; Ingram et al., 2004). It is well documented that mechanical and chemical cues relayed to VECs by their environment can alter both function and gene expression of VECs found at different tissues (Aird, 2012; Kohn et al., 2015). However, even when VECs from different origins are placed *in vitro* in the same biochemical and mechanical environment, they can still exhibit unique characteristics and behaviors that are intrinsic to the cells themselves and are not determined by differential culture or environmental conditions (Craig et al., 1998; Ostrowski et al., 2014; Stroka and Aranda-Espinoza, 2011; Ye et al., 2014). For example, the response of human umbilical cord endothelial cells (HUVEC) to changes in curvature or shear stress is completely distinct as compared to that of human brain microvascular endothelial cells (HBMEC) (Ye et al., 2014). In accordance with the differences these two cell types exhibit in response to geometrical constraints or mechanical stimuli, studies have also shown differences with respect to Lm infection. Past studies suggest that HBMEC infection by Lm depends on InlB (Greiffenberg et al., 1998), while HUVEC uptake of Lm appears to be independent of both the internalins InlA and InlB (Greiffenberg et al., 1997; Rengarajan et al., 2016). These studies are not necessarily conflicting but suggest that bacterial infection of VECs depends upon both the environment and the particular origin of VECs and therefore conclusions should refer to the exact cell type and not to endothelial cells in general. Since many pathological processes occur at the microvascular level of organs, we chose in this study to use HMEC-1 as our model for VECs. However, extrapolation from results obtained for HMEC-1 infection by Lm to other VECs should be performed with caution. In a similar analogy, although Lm uptake by HMEC-1 increases with increasing subendothelial stiffness that relationship could be different if HMEC-1 were infected with different bacterial pathogens. We speculate that the relationship between ECM stiffness and susceptibility to infection is likely host cell- and pathogen-specific.

In conclusion, our results provide the first strong evidence that the local mechanical environment and in particular the matrix stiffness of VECs regulates susceptibility of VECs to Lm infection. The novel mechanosensitive pathway we identified suggests that increased ECM stiffness sensed by VECs leads to enhanced FAK activity that augments the amount of vimentin exposed at the surface of VECs. Increased amount of surface vimentin in turn increases Lm adhesion and subsequent uptake by VECs. Our studies on the effect of ECM stiffness in bacterial uptake of VECs can have other significant implications, since they provide an example of how mechanotransduction can be exploited in biology and medicine facilitating the development of therapeutic interventions against bacterial infections and other diseases.

## MATERIALS AND METHODS

### Fabrication of thin two-layered polyacrylamide (PA) hydrogels on 24-well glass-bottom dishes

PA hydrogel fabrication was done as previously described with a few modifications for the specific experiments (Bastounis et al., 2011; Bastounis et al., 2014; Georges et al., 2006; Vincent et al., 2013). 24-well glass bottom plates (MatTek, P24G-1.5-13-F) were incubated for 1 h with 500 μL of 2 M NaOH. Wells were rinsed with distilled water, and 500 μL of 2% 3- Aminopropyltriethoxysilane (Sigma, 919-30-2) in 95% ethanol were added to each well for 5 min. Wells were rinsed again with water and 500 μL of 0.5% gluteraldehyde was added in each well for 30 min. Wells were rinsed with water and dried at 60°C. To prepare hydrogels of varying stiffness, mixtures containing 3-10% acrylamide (Sigma, A4058) and 0.06-0.6% bis-acrylamide (Fisher, BP1404-250) were prepared. Specifically, 0.6 kPa hydrogels contained 3% acrylamide and 0.06% bis-acrylamide, 3 kPa hydrogels contained 5% acrylamide and 0.1% bis-acrylamide, 20 kPa hydrogels contained 8% acrylamide and 0.26% bis-acrylamide, and 70 kPa hydrogels contained 10% acrylamide and 0.6% bis-acrylamide. For each stiffness, two mixtures were prepared the second of which contained 0.03% 0.1 μm diameter fluorescent beads (Invitrogen, F8803). Mixtures were then degassed for 15 min to remove oxygen, which inhibits acrylamide polymerization.

First, 0.06% APS and 0.43% TEMED were added to the solutions containing no beads to initiate polymerization and then 3.6 μL of the mixture was added at the center of each well, capped with 12-mm untreated circular glass coverslips and allowed to polymerize for 20 min. The coverslips were then lifted with a syringe needle with a small hook at its tip and then 2.4 μL of the mixture containing tracer beads was added, sandwiched again with a 12-mm untreated circular glass coverslip, gently pressed downwards by means of forceps, and allowed to polymerize for 20 min. 50 mM HEPES pH 7.5 was added in the wells, and coverslips were removed using the syringe needle and forceps. PA hydrogels were UV-sterilized for 1 h.

Hydrogels were then activated by adding 200 μL 0.5% w/v heterobifunctional crosslinker Sulfo-SANPAH (ProteoChem, c1111) in 1% DMSO and 50 mM HEPES pH 7.5 on the upper surface of the hydrogels and exposing to UV light for 10 min. After activation, the hydrogels were washed with 50 mM HEPES pH 7.5 to remove excess crosslinker and were coated with 200 μL of 0.25 mg/mL rat tail collagen I (Sigma-Aldrich, C3867) in 50 mM HEPES overnight at room temperature. Prior to seeding cells, hydrogels were incubated with cell media to allow equilibration for 30 min.

### HMEC-1 culture and infection with *Listeria monocytogenes* (Lm)

Human microvascular endothelial cells (HMEC-1, generous gift from Welch lab, UC Berkeley) were maintained in MCDB 131 medium (Fisher Sci Co, 10372-019) supplemented with 10% FBS (GemBio, 900-108), 10 ng/mL epidermal growth factor (Sigma, E9644), 1 μg/mL hydrocortisone (Sigma, H0888), and 2mM L-glutamine (Sigma, 56-85-9).

HMEC-1 were infected as previously described with the following modifications (Rengarajan et al., 2016). The day prior to infection, host cells were seeded at a density of 4 × 10^5^ cells/well for cells residing on wells of 24-well plates or at a density of 1 × 10^5^ cells/well for cells residing on wells of 48-well plates, and grown for 24 h. Lm liquid cultures were started from a plate colony, and grown overnight, shaking, at 30 °C in Brain Heart Infusion (BHI) media (BD, 211059) supplemented with 200 μg/mL streptomycin and 7.5 μg/mL chloramphenicol. The day of the infection, the O.D.600 of the overnight cultures was measured and diluted to 0.1. Samples were incubated for 2 h shaking, in the dark at 30°C in BHI media supplemented with 200 μg/mL streptomycin and 7.5 μg/mL chloramphenicol to allow the O.D. 600 to reach 0.2-0.3 and let bacteria enter the logarithmic growth phase. Bacteria were then washed 3 times with PBS to remove any soluble factors and infections were performed in normal growth media (Reed et al., 2014).

To synchronize invasion, Lm diluted in normal growth media were added on the HMEC-1 cells and samples were spun for 10 min at 200 × g prior to incubation. After 30 min of incubation at 37 °C, samples were washed 4 times in PBS and after an additional 30 min, media was replaced with media supplemented with 20 μg/mL gentamicin. Multiplicity of infection (MOI) was determined by plating bacteria at different dilutions, on BHI agar plates with 200 μg/mL streptomycin and 7.5 μg/mL chloramphenicol and measuring the number of colonies formed 2 days post-infection. A similar approach was followed when HMEC-1 were infected with *Listeria innocua* (Li), except that BHI plates with no antibiotics were used. All bacterial strains used in these studied are indicated in Table S1.

Analysis of infection via flow cytometry was performed at 7-8 h after exposure, unless otherwise stated. For drug exposure experiments, unless otherwise indicated, media was removed from the cells and replaced with media containing either the drug or vehicle control 1 h prior to infection or 30 min prior to infection for the case of withaferin. Cells were cultured in the drug-containing media for 1 h after infection. 1 h post-infection cells were washed four times and drug-free gentamicin-containing media was added to the cells.

### Antibodies and reagents

Hoechst (Thermofisher, D1306) was dissolved at 1 mg/mL in dimethyl formamide (DMSO) and used at 1:1000. Drugs were dissolved in DMSO (Sigma, D2650) at the stock concentrations indicated: 10 mM FAK inhibitor-14 (FAK-14) (Tocris Bioscience, 3414), 100 mM PF573228 (Tocris Bioscience, 3239), 20 mM SGX 523 (Tocris Bioscience 5356), 42 mM withaferin (Sigma-Aldrich, W4394). Primary antibody used for staining of extracellular Lm was rabbit polyclonal anti-Listeria genus specific antibody (ABCAM, ab35132). Primary antibody used for staining Li was goat polyclonal BacTrace *anti-Listeria* genus specific antibody (SeraCare Life Sciences Inc., 01-90-90). For incubation of cells with vimentin antibodies prior to infection or immunostaining of surface vimentin, rabbit polyclonal H-84 vimentin antibody (Santa Cruz Biotechnologies, sc-5565) was used. For isotype controls, cells were incubated with rabbit IgG (Sigma, I5006). For western blotting the following antibodies were used: rabbit polyclonal phospho-FAK Tyr397 (ThermoFisher, 44-624G), mouse monoclonal anti-FAK (EMD Millipore, 05-182), rabbit monoclonal anti-Met (cMet) antibody (Abcam, ab51067), anti-GAPDH antibody (Cell signaling, 140C10).

### Flow cytometry of HMEC-1 cells infected with Lm

7-8 h post-infection, infected HMEC-1 cells were detached from the substrate by incubating them for 10 min with a mixture of 200 μL of 0.25% trypsin-EDTA and 0.05% collagenase (Sigma, C0130). Solutions in each well were pipetted up and down 6 times to ensure single cell suspensions, and 200 μL of complete media was added to inactivate trypsin in each well. Solutions were transferred into 5 mL polystyrene tubes with a 35-μm cell strainer cap (Falcon, 352235) and then samples were immediately analyzed by flow cytometry on the Scanford FACScan analyzer (Custom Stanford and Cytek upgraded FACScan). 10,000-20,000 cells were analyzed per each replicate. To ensure analysis of single cells, the bulk of the distribution of cell counts was gated using the forward versus side scatter plot. This gating strategy ensures that single cells are analyzed and debris or cell doublets or triplets are eliminated from the analysis. A second gating step was then performed to exclude cells that autofluoresce by measuring the fluorescence of control-uninfected cells and gating the population of infected cells so to exclude autofluorescence.

### Immunostaining of extracellular adherent bacteria

HMEC-1 cells residing on either PA hydrogel substrates or glass collagen I-coated coverslips were infected as described above with a Lm strain that constitutively expresses GFP. 20 min post-infection, 1 μg/mL Hoechst (Thermofisher, D1306) was added in each well to stain the cells’ nuclei. 30 min post-infection cells were washed 4 times in PBS and fixed with a non-permeabilizing fixative for differential immunostaining for 20 min at room temperature (Yam and Theriot, 2004). Fixative contained 0.32 M sucrose, 10mM MES pH 6.1, 138 mM KCl, 3 mM MgCl_2_, 2 mM EGTA and 4% formaldehyde EM grade. Following a wash with PBS, samples were blocked for 30 min with 5% BSA in PBS and then incubated with anti-Lm primary antibody (Abcam ab35132) diluted 1:100 in PBS containing 2% BSA for 1 h. Samples were washed in PBS 3 times and then incubated with Alexa Fluor 546 goat anti-rabbit secondary antibody (Invitrogen A-11035) diluted 1:250 in PBS containing 2% BSA for 1 h. Samples were washed 3 times in PBS and stored in 1 mL PBS for imaging. N > 1000 cells were analyzed per condition. For imaging, we used an inverted Nikon Diaphot 200 with a CCD camera (Andor Technologies) and a 40X air Plan Fluor NA 0.60 or a 100X oil objective. The microscope was controlled by the MicroManager software package (Edelstein et al., 2014). For differential immunostaining, all “green” bacteria associated with individual cells were counted as adherent; bacteria that were both “green” and ȁred” (due to antibody binding) were counted as non-internalized. Nuclei number was identified running a custom-made script in MATLAB (Mathworks Inc.) and the CellC software was used for enumeration of the bacteria (Selinummi et al., 2005). For characterization of total bacteria adhering to host cells (Lm or Li), same procedure was followed with the exception that cells were permeabilized for 5 min in 0.2% Triton-X in PBS. Primary antibody used was goat polyclonal BacTrace anti-Listeria genus specific antibody.

### Immunostaining of surface vimentin

HMEC-1 cells residing on glass collagen I-coated coverslips were incubated at 37°C for 30 min with H-84 anti-vimentin antibody in 1:10 dilution in media. Cells were then washed three times in PBS and fixed in 4% formaldehyde (EM grade) in PBS for 10 min at room temperature. A blocking step with 5% FBS in PBS was followed for 30 min and then cells were incubated with secondary goat anti-rabbit AlexaFluor-546 antibody diluted 1:250 in PBS containing 2% BSA for 1 h. For negative controls to ensure label is specific to primary antibody, cells with no primary antibody were fixed and incubated for 1 h with secondary goat anti-rabbit AlexaFluor-546 antibody. As an additional control, cells were also fixed, permeabilized and stained for intracellular vimentin. For imaging, we used an inverted Nikon Diaphot 200 with a CCD camera (Andor Technologies) and a 100X oil objective.

### HMEC-1 cell transfection with siRNA

For each well of a 24-well plate, 6 × 10^4^ HMEC-1 cells suspended in serum free media were reverse-transfected with siRNAs at 20 nM final concentration using 0.25 μL lipofectamine RNAiMAX (Invitrogen 13778075). The transfection mix was replaced by full media 8 h later. Synthetic siRNA pools (including 4 distinct siRNA sequences for each gene) to target vimentin, Met and FAK were purchased from Dharmacon (Table S2). HMEC-1 cells were treated with control (non-targeting, siGLO and Kif11) or experimental siRNA in accordance with the manufacturer instructions. Specifically, to demonstrate that transfection performed was sufficient to get siRNAs into the cells, we transfected cells with synthetic siRNA, siGLO, that makes cell exposed to it fluorescent 24 h post-transfection (Table S2). In addition to track a cell cycle phenotype to verify that knockdown has occurred with our protocol, we transfected cells with siKif11 that results in substantial cell death of transfected cells, approximately 24-48 h post-transfection and can be verified on the TC microscope. Bacterial infections were performed approximately 72 h after transfection

### RT-qPCR

HMEC-1 cells were treated with control or experimental siRNA as described above. mRNA was harvested using the RNeasy Micro Kit (Qiagen, 74004) and cDNA was prepared using the Superscript III First-strand Synthesis SuperMix (ThermoFisher, 18080–400). qPCR was performed using TaqMan PreAmp Master (Thermofisher, 4331182). Genes of interest were amplified using primers Hs00958113_g1 for Vimentin (ThermoFisher, 4331182) and Hs99999905_m1 for GAPDH (ThermoFisher, 4333764) on a StepOnePlus Real-Time PCR System. Normalized relative quantity (NRQ) and error were calculated as previously described [92]. GAPDH was used as control gene.

### Western blotting of FAK and Met for HMEC-1 lysates coming from cells residing on different substrates

For assessing FAK and Met phosphorylation and expression levels, cells were seeded on different substrates for 24 h, and then lysed with a buffer containing 1% Nonidet P-40, 0.5% sodium deoxycholate, and a protease inhibitor mixture (PMSF, leupeptin, aprotinin, and sodium orthovanadate). The total cell lysate was separated by SDS/PAGE (10% running, 4% stacking) and transferred onto a nitrocellulose membrane (Immobilon P, 0.45-μm pore size). The membrane was then incubated with the designated antibodies. Immunodetection was performed using the Western-Light chemiluminescent detection system (Applied Biosystems).

### Cell surface protein isolation

The isolation of cell surface proteins was adapted according to (Karhemo et al., 2012) and (Roesli et al.). Cells were plated on 10 × 15 cm cell culture dishes until reaching confluency for each condition. Cells from 10 dishes were incubated with vehicle control (DMSO) containing media and cells from other 10 dishes with 2 μM PF537228 FAK inhibitor for 1 h. Cells were then washed three times with DPBS+ (PBS, 0.901 mM CaCl2, 0.492 mM MgCl_2_, 2.667 mM KCl, 1.47 mM KH2PO4, 137.931 mM NaCl, 8.060 mM Na2HPO4-7H2O, 5.555 mM D-Glucose, 0.327 mM Sodium Pyruvate), and incubated for 30 min with 0.5 mM EZ-Link Sulfo-NHS-SS-Biotin (APExBIO, 21331) in DPBS+ at 4°C. To quench any non-reacted biotinylation, 1 M Tris pH 8.0 at a final concentration of 50 mM was added on each plate for 10 min at 4°C. The solution was then discarded and plates were rinsed twice in 1X TBS (50 mM Tris-Cl pH 7.2, 150 mM NaCl). 2 ml of TBS pH 7.2 with 1X protease inhibitor cocktail (1 mM EDTA, 10 μM E-64, 2 μg/ml leupeptin, 15 μg/ml benzamidine, 0.1 mM PMSF) and with 1X phosphatase inhibitor cocktail (1 mM Na_3_VO_4_, 1 mM NaF) were added to each dish and cells were scraped into 40 ml falcon tubes. Tubes were spun at 400 × g for 5 min and the supernatant was discarded. 10 ml of lysis buffer (50 mM Tris-HCL pH 8.5, 150 mM NaCl, 2% NP-40, 0.25% Deoxycholate, 2x protease inhibitor cocktail, 1x osphatase inhibitor cocktail) were added to each single pellet obtained from 10 × 15 cm plates, and cells were passed through a 25 G needle attached to a 10-mL syringe 3 times. Lysates were then sonicated for 5 min in a bioruptor with 30 sec ON/OFF cycle. 1 mM MgCl_2_ and 50 U/ml benzonase (Sigma, 9025-65-4) were added to the samples that were then incubated at 4°C for 2 h on a rotator. EDTA to a final concentration of 5 mM was added to each sample and samples were spun in an ultracentrifuge (Rotor type 70.1 Ti; 33,000 rpm) at 100,000x g for 60 min at 4°C. Supernatants were then transferred into fresh 15 mL Falcon tubes and protein concentration was estimated by the BCA protein assay (Thermo Fisher Scientific, Pierce Rockford). Equal amounts of protein (~5 mg) from each extract were used for cell surface protein isolation.

Streptavidin agarose beads (Pierce, 20347) were washed three times with lysis buffer.50 μL per 5 mg/lysate were added for 2 h or overnight at 4 °C on a rotator. Samples were then transferred to 2 ml gravity flow columns (BioRad) pre-equilibrated with 2 ml wash buffer I (50 mM Tris-HCL pH 8.5, 150 mM NaCl, 2% NP-40, 0.25% Deoxycholate, 0.5% SDS). The flow through was collected and stored at −80 °C. Columns were washed once with 2 mL lysis buffer (excluding protease and phosphatase inhibitors), then with 2 mL wash buffer I, followed by a 2 ml wash with wash buffer II (50 mM Tris-HCL pH 8.5, 300 mM NaCl, 2% NP-40, 0.25% Deoxycholate, 0.5% SDS) and then again with 2 mL wash buffer I. Beads were then transferred to a fresh 1.5 mL Eppendorf tube and 3x volume of elution buffer (100 mM Tris-HCl pH 8.5, 2% SDS, 100 mM DTT) was added and samples were incubated at 37 °C for 1 h on a thermomixer. Beads were then spun down and supernatant was collected and 5% aliquoted for gel-analysis (2.5% elution for Western Blot and 2.5% elution for Silver staining of 1D-PAGE gels) and the rest was put in a fresh 1.5 mL Eppendorf tube and stored at −80°C.

### 2D SDS-PAGE electrophoresis and staining

Two-dimensional electrophoresis was performed according to the carrier ampholyte method of isoelectric focusing (Burgess-Cassler et al., 1989; O’Farrell, 1975) by Kendrick Labs, Inc. (Madison, WI) as follows: Isoelectric focusing was carried out in a glass tube of inner diameter 2.3 mm using 2% pH 3-10 Isodalt Servalytes (Serva, Heidelberg, Germany) for 9600 volt-h. 1 μg of an IEF internal standard, tropomyosin, was added to the sample. This protein migrates as a doublet with lower polypeptide spot of MW 33,000 and pI 5.2. The enclosed tube gel pH gradient plot for this set of Servalytes was determined with a surface pH electrode. After equilibration for 10 min in Buffer ‘O’ (10% glycerol, 50 mM dithiothreitol, 2.3% SDS and 0.0625 M Tris, pH 6.8), each tube gel was sealed to the top of a stacking gel that overlaid a 10% acrylamide slab gel (0.75 mm thick). SDS slab gel electrophoresis was carried out for about 4 h at 15 mA/gel. The following proteins (Sigma Chemical Co., St. Louis, MO and EMD Millipore, Billerica, MA) were used as molecular weight standards: myosin (220,000), phosphorylase A (94.000),catalase (60,000), actin (43,000), carbonic anhydrase (29,000) and lysozyme (14.000). These standards appear along the basic edge of the silver-stained (Oakley et al., 1980) 10% acrylamide slab gel. The gels were dried between sheets of cellophane with the acid edge to the left.

### Manual comparisons of patterns of silver-stained 2D gels

Each gel for comparison was overlaid with a transparent sheet for labeling polypeptide spot differences without marking the original gel. Two experienced analysts manually compared protein patterns and polypeptide spots that were unique to or differed between controls and PF537228 samples were outlined.

### Protein digestion and peptide extraction

Proteins that were separated by SDS-PAGE/2D-PAGE and stained by Coomassie dye were excised, washed and the proteins from the gel were treated according to published protocols (Darie et al., 2011; Shevchenko et al., 1996; Sokolowska et al., 2012c). Briefly, the gel pieces were washed in high purity, high performance liquid chromatography HPLC grade water, dehydrated and cut into small pieces and de-stained by incubating in 50 mM ammonium bicarbonate, 50 mM ammonium bicarbonate/50% acetonitrile, and 100% acetonitrile under moderate shaking, followed by drying in a speed-vac concentrator. The gel bands were then rehydrated with 50 mM ammonium bicarbonate. The procedure was repeated twice. The gel bands were then rehydrated in 50 mM ammonium bicarbonate containing 10 mM DTT and incubated at 56°C for 45 min. The DTT solution was then replaced by 50 mM ammonium bicarbonate containing 100 mM Iodoacetamide for 45 min in the dark, with occasional vortexing. The gel pieces were then re-incubated in 50 mM ammonium bicarbonate/50% acetonitrile, and 100% acetonitrile under moderate shaking, followed by drying in speed-vac concentrator. The dry gel pieces were then rehydrated using 50 mM ammonium bicarbonate containing 10 ng/μL trypsin and incubated overnight at 37°C under low shaking. The resulting peptides were extracted twice with 5% formic acid/50 mM ammonium bicarbonate/50% acetonitrile and once with 100% acetonitrile under moderate shaking. Peptide mixture was then dried in a speed-vac, solubilized in 20 μL of 0.1% formic acid/2% acetonitrile.

### Liquid chromatography and tandem mass spectrometry (LC-MS/MS)

The peptide mixture was analyzed by reverse phase liquid chromatography (LC) and MS (LC-MS/MS) using a NanoAcuity UPLC (Micromass/Waters, Milford, MA) coupled to a Q-TOF Ultima API MS (Micromass/Waters, Milford, MA), according to published procedures (Darie et al., 2011; Sokolowska et al., 2012a; Sokolowska et al., 2012b; Spellman et al., 2008). Briefly, the peptides were loaded onto a 100 μm × 10 mm NanoAquity BEH130 C18 1.7 μm UPLC column (Waters, Milford, MA) and eluted over a 150 min gradient of 2-80% organic solvent (ACN containing 0.1% FA) at a flow rate of 400 nL/min. The aqueous solvent was 0.1% FA in HPLC water. The column was coupled to a Picotip Emitter Silicatip nano-electrospray needle (New Objective, Woburn, MA). MS data acquisition involved survey MS scans and automatic data dependent analysis (DDA) of the top three ions with the highest intensity ions with the charge of 2+, 3+ or 4+ ions. The MS/MS was triggered when the MS signal intensity exceeded 10 counts/sec. In survey MS scans, the three most intense peaks were selected for collision-induced dissociation (CID) and fragmented until the total MS/MS ion counts reached 10,000 or for up to 6 sec each. The entire procedure used was previously described (Darie et al., 2011; Sokolowska et al., 2012a; Sokolowska et al., 2012b). Calibration was performed for both precursor and product ions using 1 pmol GluFib (Glu1-Fibrinopeptide B) standard peptide with the sequence EGVNDNEEGFFSAR and the monoisotopic doubly-charged peak with m/z of 785.84.

### LC-MS/MS data processing and protein identification

The raw data were processed using ProteinLynx Global Server (PLGS, version 2.4) software as previously described (Sokolowska et al., 2012a). The following parameters were used: background subtraction of polynomial order 5 adaptive with a threshold of 30%, two smoothings with a window of three channels in Savitzky-Golay mode and centroid calculation of top 80% of peaks based on a minimum peak width of 4 channels at half height. The resulting pkl files were submitted for database search and protein identification to the public Mascot database search (www.matrixscience.com, Matrix Science, London, UK) using the following parameters: databases from NCBI (bacteria), parent mass error of 1.3 Da, product ion error of 0.8 Da, enzyme used: trypsin, one missed cleavage, propionamide as cysteine fixed modification and Methionine oxidized as variable modification. To identify false negative results, we used additional parameters such as different databases or organisms, a narrower error window for the parent mass error (1.2 and then 0.2 Da) and for the product ion error (0.6 Da), and up to two missed cleavage sites for trypsin. In addition, the pkl files were also searched against in-house PLGS database version 2.4 (www.waters.com) using searching parameters similar to the ones used for Mascot search. The Mascot and PLGS database search provided a list of proteins for each gel band. To eliminate false positive results, for the proteins identified by either one peptide or a mascot score lower than 25, we verified the MS/MS spectra that led to identification of a protein.

### Atomic force microscopy force-distance measurements for hydrogel stiffness characterization

AFM force-distance beating were performed on PA hydrogel samples in 50mM HEPES pH 7.5 buffer with a Park NX-10 AFM (Park Systems, Santa Clara, CA) using commercial silicon nitride cantilevers CP-PNP-SiO with a sphere tip (sQube, 0.08 N/m stiffness, sphere radius ~ 1 micron) and gold coating on the reflective side. Temperature has been kept at 37 °C throughout the experiment. Tip calibration curves were performed on glass substrate surface, which was considered infinitely hard for the soft tips used. Two approach-withdraw cycles were performed per cell. Data analysis of FD curves and calculation of Young’s modulus was performed using XEI software (Park Systems, Santa Clara, CA) and SPIP software (Image Metrology, Hørsholm, Denmark).

### Visualization and comparison of distributions

Distributions were visualized by using boxplots with the features indicated below. Bold black line parallel to the x-axis indicates the median (second quartile) of the distribution and boxplot’s notched section shows the 95% confidence interval around the median (Wilcoxon-Mann-Whitney nonparametric test). Regular black lines extended vertically below and above the medians represent the first and third quartiles of the distribution. Lines extending vertically from the boxplot (whiskers) represent the lowest datum within 1.5 IQR (interquartile range) of the lower quartile and the highest datum within 1.5 IQR of the upper quartile. Data points beyond the whiskers are considered outliers and displayed as circles. To assess whether the differences between the medians of two distributions are significant we run a nonparametric Wilcoxon rank sum test since the distributions are not normal. One or two asterisks denote statistically significant differences between the medians of two distributions (<0.05 or <0.01, respectively).

## AUTHOR CONTRIBUTIONS

E.E.B. and J.A.T. conceived the project and designed the experiments. E.E.B. performed experiments and analyzed data. Y.Y. performed experiments. E.E.B. and J.A.T. wrote the paper.

## ACKNOWLEDGEMENTS

Our thanks to M. Footer, R. Lamason, M. Rengaranjan, G. Skariah and members of the Theriot Lab for discussions and experimental support. This work was supported by NIH R01AI036929 (J.A.T), HHMI (J.A.T) and the American Heart Association (E.E.B. and Y.Y.). Flow cytometry was performed at the Stanford Shared FACS Facility and RT-qPCR was performed at the Stanford PAN Facility.

## SUPPORTING INFORMATION CAPTIONS

**Figure S1.**
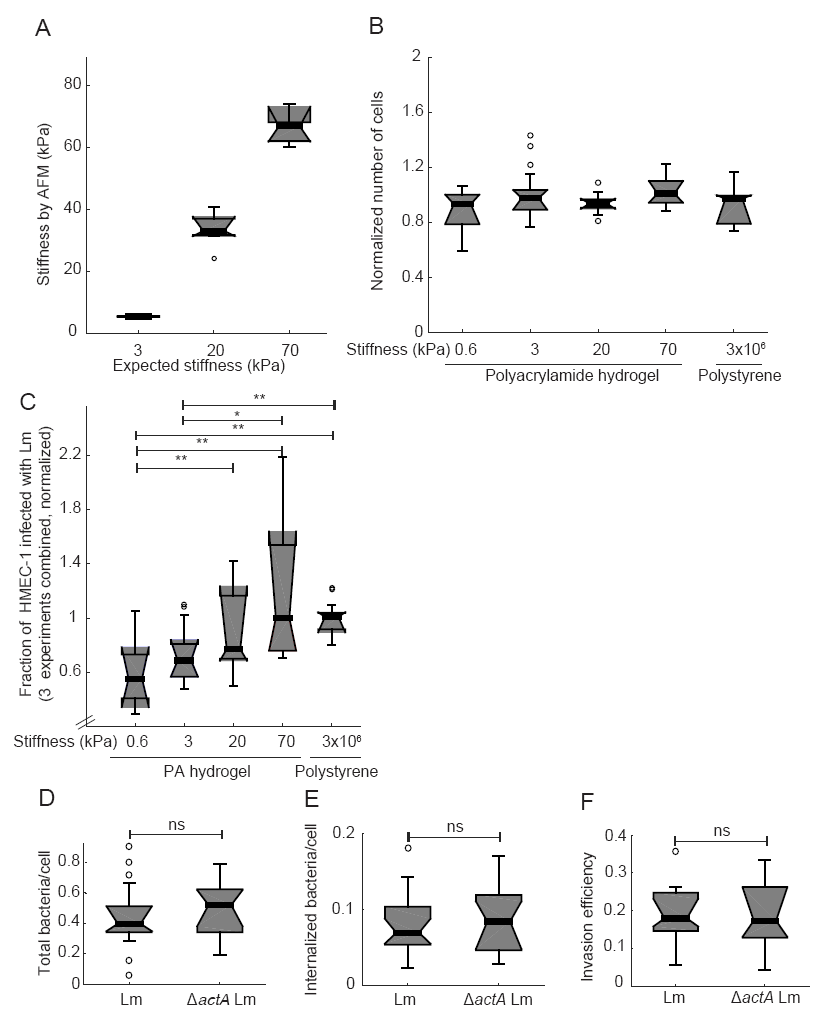
Lm infection of VECs residing on hydrogels of varying stiffness. (**A**)Young’s moduli of hydrogels. Plot showing the expected Young’s modulus of the hydrogels given the amount of acrylamide and bis-acrylamide used versus the Young’s modulus measured through AFM (N = 6). Stiffness of the 0.6 kPa could not be measured, as the hydrogels were very soft and adhered to the AFM tip.(B) Fraction of cells passed through the flow cell in 20 s for HMEC-1 residing on varying stiffness substrates. HMEC-1 were seeded on 0.6 kPa, 3 kPa, 20 kPa and 70 kPa PA hydrogels as well as TC polystyrene substrates for 24 h at a starting concentration of 4 × 10^5^ cells per well for 24 h. Cells were then detached from their matrix and number of cells per well passed through the flow cell in 20 s was measured using a flow cytometry for cells coming from N=5-6 wells for each substrate stiffness. Results from 3 independent experiments were combined and for each experiment values were normalized relative to the mean number of cells passed through the flow cell for cells residing on TC polystyrene substrates. No significant differences were detected. (**C**) Boxplots of normalized fraction of HMEC-1 infected with *ΔactA* Lm for cells residing on PA hydrogels of varying stiffness or TC polystyrene. Stiffness (kPa) of each substrate is indicated. Boxplots refer to data from 3 independent experiments, with N = 5-6 replicates per experiment. For each experiment values have been normalized relative to the mean infection level of cells residing on polystyrene substrates. (**D-F**) HMEC-1 residing on collagen I-coated glass substrates were infected with Lm (constitutively expressing GFP) or *ΔatcA* Lm at an MOI of 2.7. 30 min post-infection samples were fixed, immunostained and infection was analyzed by microscopy followed by image processing. Boxplots showing: (**D**) total bacteria per cell; (**E**) internalized bacteria per cell; (**F**) invasion efficiency (ratio of internalized bacteria to total bacteria). For each condition, 500-600 cells were analyzed in total. Representative data come from 1 of 2 independent experiments. Circles represent outliers, and the boxplots’ notched sections show the 95% confidence interval around the median (Wilcoxon-Mann-Whitney test, for details about boxplots see Materials and Methods). One or two asterisks denote statistically significant differences between the medians of two distributions (<0.05 or <0.01, respectively; Wilcoxon rank sum test).

**Figure S2.**
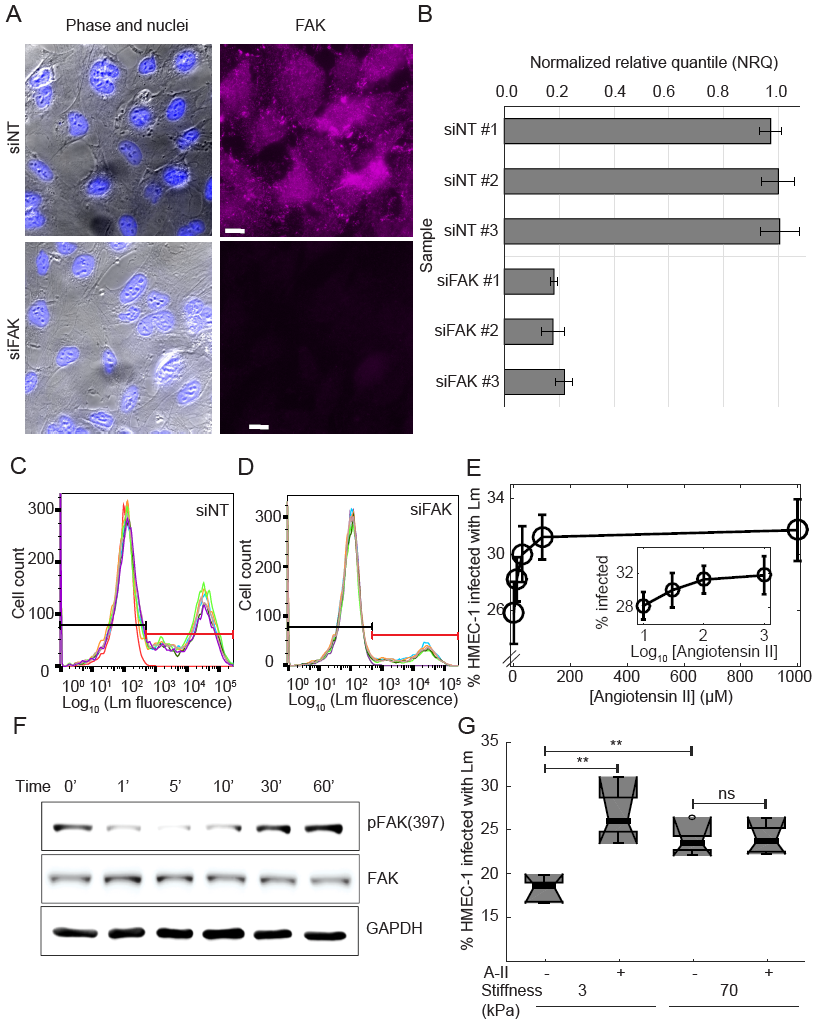
FAK-dependent decrease in bacterial uptake. (**A**) HMEC-1 treated with 20 nM of non-targeting siRNA (siNT, first row) or FAK siRNA (siFAK, second row) were stained for FAK using an anti-FAK antibody. Representative phase image of cells (left column), superimposed with the image of the nuclei (blue) and anti-FAK antibody fluorescence (purple, right column) are shown. Scale bar shown in white is 23 μm. (**B**) Relative with respect to GAPDH gene expression of FAK obtained by RT-qPCR. The levels relative levels of expression in each siRNA treated sample (#1-#3) are expressed relative to the control siNT treated sample #2 (normalized relative quantity, NRQ). N=3 replicates are shown for each group treated with either siNT or siVIM. (C-D) Histograms of the logarithm of Lm fluorescence intensity per cell for HMEC-1 treated with 20 nM of non-targeting siRNA (siNT) (C) or FAK siRNA (siFAK) (D). HMEC-1 were infected with *ΔactA* Lm (actAp::mTagRFP) and infection was analyzed by flow cytometry, 7-8 h after infection. MOI is 45. Histograms for N=6 replicates are shown in different colors. Control uninfected cells’ histogram is shown in purple. Based on the autofluorescence of the control group a gate is defined (see black and red lines) showing what is considered uninfected (left, black line) and infected (right, red line). (E) Increase in bacterial uptake by Angiotensin II. Angiotensin II or vehicle control was added 2 h before addition of bacteria to HMEC-1 residing on polystyrene substrates. Percentage of HMEC-1 infected with *ΔactA* Lm (actAp::mTagRFP) as a function of inhibitor concentration (mean +/− SD, N = 4 replicates). Infection was analyzed by flow cytometry, 7-8 h after infection. MOI is 70. (**F**) Western blots from whole HMEC-1 lysates showing expression of phosphorylated FAK (Tyr397) and total FAK for cells residing on TC polystyrene substrates and treated for varying amount of time with 100 nM Angiotensin-II. In each Western blot, equal quantities of protein were loaded and equal loading was confirmed in relation to glyceraldehyde 3-phosphate dehydrogenase (GAPDH) expression. The Western blots shown are representative of 3 independent experiments. (**G**) Boxplots of percentage of HMEC-1 infected with *ΔactA* Lm (actAp::mTagRFP) for cells residing on soft (3 kPa) or stiff (70 kPa) hydrogels and pre-treated for 2 h either with vehicle control or 100 nM Angiotensin-II (N=6 replicates). MOI is 62.5. Two asterisks denote statistically significant differences between the medians of two distributions (<0.01; Wilcoxon rank sum test).

**Figure S3.**
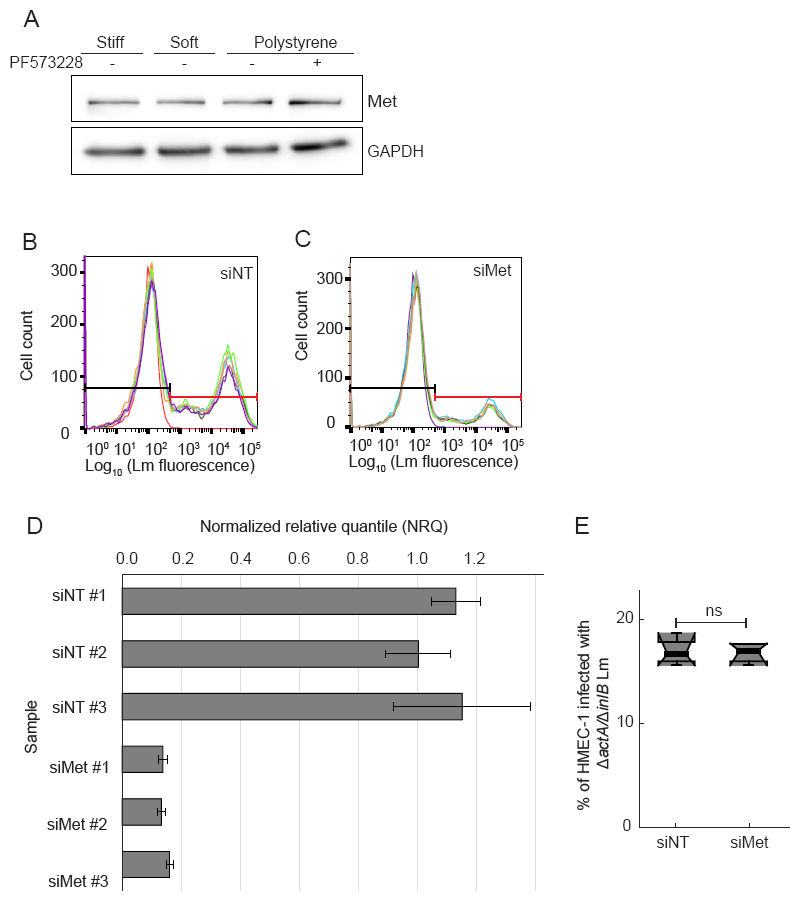
Met-dependent decrease in bacterial uptake. (**A**) Western blot from whole HMEC-1 lysates showing expression of Met for cells seeded on soft (3 kPa), stiff (70 kPa) and TC polystyrene substrates treated or not with 2 μM PF537228 FAK inhibitor. In each case, equal quantities of protein were loaded and equal loading was also confirmed in relation to glyceraldehyde 3-phosphate dehydrogenase (GAPDH) expression. (**B-C**) Histograms of the logarithm of Lm fluorescence intensity per cell for HMEC-1 treated with 20 nM of non-targeting siRNA (siNT) (**C**) or Met siRNA (siMet) (**D**). HMEC-1 were infected with *ΔactA* Lm (actAp::mTagRFP) and infection was analyzed by flow cytometry, 7-8 h after infection. MOI is 45. Histograms for N=6 replicates are shown in different colors. Control uninfected cells’ histogram is shown in purple. Based on the autofluorescence of the control group a gate is defined (see black and red lines) showing what is considered uninfected (left, black line) and infected (right, red line). (**D**) Relative with respect to GAPDH gene expression of Met obtained by RT-qPCR. The levels relative levels of expression in each siRNA treated sample (#1-#3) are expressed relative to the control siNT treated sample #2 (normalized relative quantity, NRQ). N=3 replicates are shown for each group treated with either siNT or siVIM. (E) Boxplots of percentage of HMEC-1 infected with *ΔactA/ΔinlB* Lm (actAp::mTagRFP) for cells HMEC-1 treated with siNT or siMet (N=4 replicates). MOI is 60.

**Figure S4.**
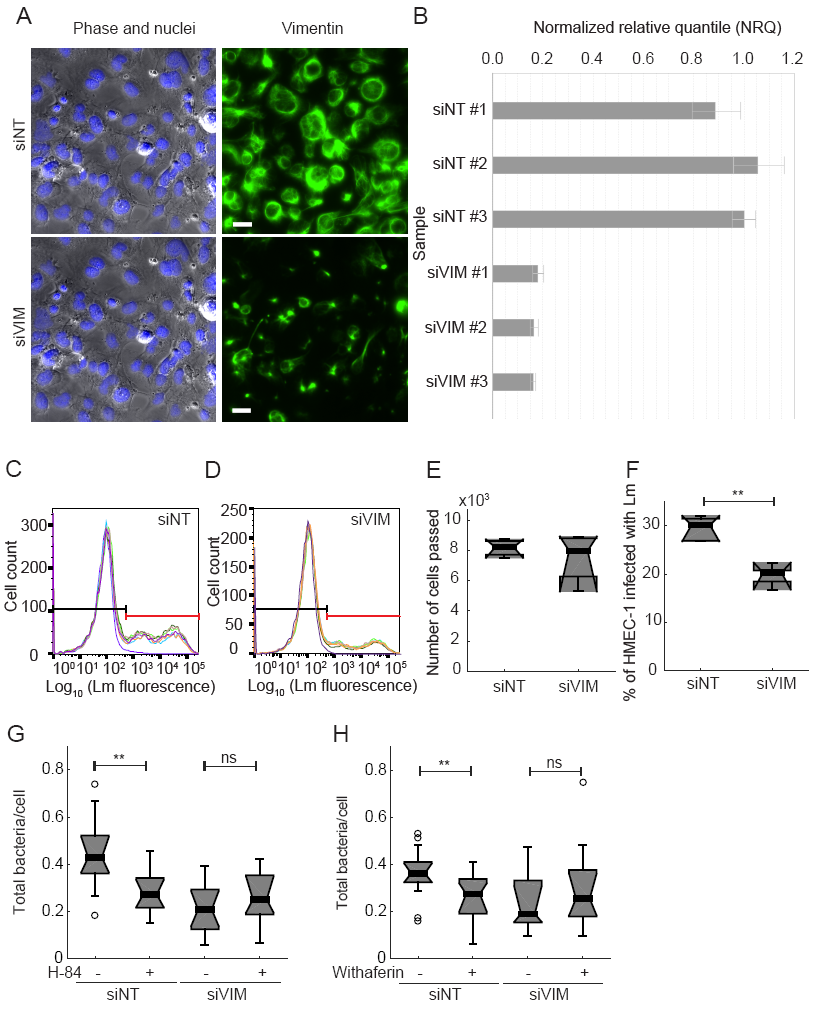
Knocking down vimentin decreases Lm uptake. (**A**) HMEC-1 treated with 20 nM of non-targeting siRNA (siNT, first row) or VIM siRNA (siVIM, second row) were stained for VIM using the H-84 anti-vimentin antibody. Representative phase image of cells (Phase, left column), superimposed with the image of the nuclei (blue) and anti-vimentin antibody fluorescence (green, right column) are shown. Scale bar shown in white is 35 μm. (**B**) Relative with respect to GAPDH gene expression of vimentin obtained by RT-qPCR. The levels relative levels of expression in each siRNA treated sample (#1-#3) are expressed relative to the control siNT treated sample #2 (normalized relative quantity, NRQ). N=3 replicates are shown for each group treated with either siNT or siVIM (C-D) Histograms of the logarithm of Lm fluorescence intensity per cell for HMEC-1 treated with 20 nM of non-targeting siRNA (siNT) (**C**) or vimentin siRNA (siVIM) (**D**). HMEC-1 were infected with *ΔactA* Lm (actAp::mTagRFP) and infection was analyzed by flow cytometry, 7-8 h after infection. MOI is 34. Histograms for N=6 replicates are shown in different colors. Control uninfected cells’ histogram is shown in purple. Based on the autofluorescence of the control group a gate is defined (see black and red lines) showing what is considered uninfected (left, black line) and infected (right, red line). (**E**) Number of HMEC-1 passed through the flow cell in 20 s, for N = 4 samples coming from wells treated with either siNT or siVIM. (**F**) Boxplots of percentage of HMEC-1 infected with *ΔactA* Lm for the data shown in panels C-D. One or two asterisks denote statistically significant differences between the medians of two distributions (<0.05 or <0.01, respectively; Wilcoxon rank sum test). (**F**) HMEC-1 residing on collagen I-coated glass substrates, treated with siNT or siVIM were also blocked for 1 h with anti-vimentin antibodies prior to infection. Cells were infected with Lm (constitutively expressing GFP) at an MOI of 1.25. 30 min post-infection samples were fixed, immunostained and infection was analyzed by microscopy followed by image processing. Boxplots shows total bacteria per cell for an average of N=650 cells analyzed per condition (**F**) HMEC-1 residing on collagen I-coated glass substrates, treated with siNT or siVIM were also treated for 30 min with vehicle control or 5 μM withaferin prior to infection. Cells were infected with Lm (constitutively expressing GFP) at an MOI of 1.25. 30 min post-infection samples were fixed, immunostained and infection was analyzed by microscopy followed by image processing. Boxplots shows total

**Figure S5.**
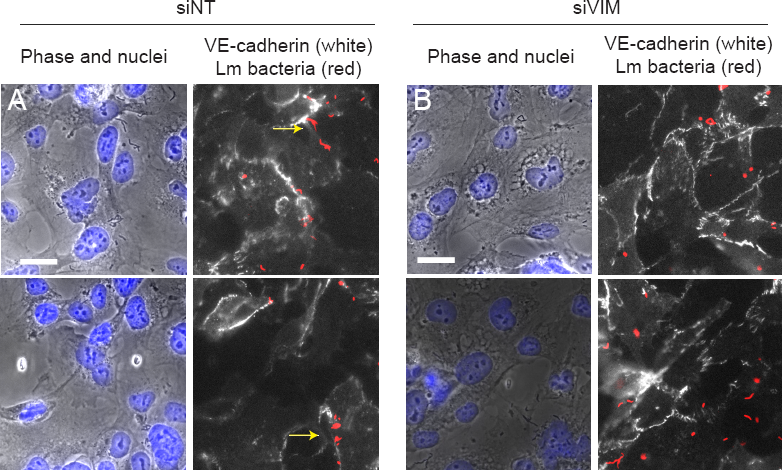
Fewer Lm localize at cell-cell junctions for siVIM treated HMEC-1. (**A-B**) HMEC-1 treated with 20 nM of non-targeting siRNA (siNT, A) or vimentin siRNA (siVIM, B) were infected with Lm at an MOI of 3. 30 min post-infection samples were fixed, stained for VE-cadherin that localizes all throughout the cell-cell junctions using an anti-VE-cadherin antibody. Representative phase image of cells (left column), superimposed with the image of the nuclei (blue) and anti-VE-cadherin antibody fluorescence (white, right column) superimposed with fluorescence of Lm (red). Scale bar is 23 μm. Two representative examples are shown for each condition. Yellow arrows point at cell-cell junctions were bacteria are co-localized.

## SUPPLEMENTARY TABLES

**Supplementary Table 1.** Bacterial strains used in this study.

| Name and source | Species | Genotype/Description |
| --- | --- | --- |
| JAT983 (100) | *L. monocytogenes* 10403S | LLO<sup>G486D</sup><br>actAp::mTagRFP |
| JAT985 (100) | *L. monocytogenes* 10403S | LLO<sup>G486D</sup> $\Delta actA$<br>actAp::mTagRFP |
| JAT1045 (100) | *L. monocytogenes* 10403S | LLO<sup>G486D</sup><br>Constitutive sGFP |
| JAT1046 (100) | *L. monocytogenes* 10403S | LLO<sup>G486D</sup> $\Delta actA$<br>Constitutive sGFP |
| JAT1063 (100) | *L. monocytogenes* 10403S | LLO<sup>G486D</sup> $\Delta actA/\Delta inlB$<br>actAp::mTagRFP |
| JAT1060 (100) | *L. monocytogenes* 10403S | LLO<sup>G486D</sup> $\Delta inlB$<br>Constitutive sGFP |
| JAT1051 (100) | *L. monocytogenes* 10403S | LLO<sup>G486D</sup> $\Delta inlA$<br>actAp::mTagRFP |
| JAT1062 (100) | *L. monocytogenes* 10403S | LLO<sup>G486D</sup> $\Delta inlB$<br>actAp::mTagRFP |
| JAT1115 (100) | *L. monocytogenes* 10403S | LLO<sup>G486D</sup> $\Delta inlF$<br>actAp::mTagRFP |
| JAT638 (DP-L392) (85) | *L. innocua* |  |

**Supplementary Table 2.** Synthetic siRNA pools used in this study.

| Accession number | Gene (Protein) | Product Name | Catalog Number |
| --- | --- | --- | --- |
|  | Negative control | ON-TARGETplus<br>Non-Targeting Pool | D-001810-10 |
| NM_000942 | siGLO | ON-TARGETplus<br>siGLO Control<br>Reagent | D-001610-01 |
| NM_004523.3 | Kif11 | ON-TARGETplus<br>Human KIF11<br>(3832) siRNA | L-003317-00-0005 |
| NM_003380.3 | VIM | ON-TARGETplus<br>Human VIM | L-003551-00-0005 |
| NM_000245.3 | MET | ON-TARGETplus<br>Human MET | L-003156-00-0005 |
| NM_001199649.1 | PTK2 (FAK) | ON-TARGETplus<br>Human PTK2 | L-003164-00-0005 |

